# Dissecting the co-transcriptome landscape of plants and microbiota members

**DOI:** 10.1101/2021.04.25.440543

**Authors:** Tatsuya Nobori, Yu Cao, Frederickson Entila, Eik Dahms, Yayoi Tsuda, Ruben Garrido-Oter, Kenichi Tsuda

**Affiliations:** State Key Laboratory of Agricultural Microbiology, Hubei Hongshan Laboratory, Hubei Key Lab of Plant Pathology, College of Plant Science and Technology, Huazhong Agricultural University, Wuhan 430070, China.; Shenzhen Institute of Nutrition and Health, Huazhong Agricultural University, Wuhan 430070, China.; Shenzhen Branch, Guangdong Laboratory of Lingnan Modern Agriculture, Genome Analysis Laboratory of the Ministry of Agriculture and Rural Affairs, Agricultural Genomics Institute at Shenzhen, Chinese Academy of Agricultural Sciences, Shenzhen, Guangdong 518120, China; Department of Plant Microbe Interactions, Max Planck Institute for Plant Breeding Research, Carl-von-Linne-Weg 10, Cologne 50829, Germany.; Salk Institute for Biological Studies, La Jolla, CA 92037, USA.; Cluster of Excellence on Plant Sciences, 40225 Düsseldorf, Germany.

## Abstract

Interactions between plants and neighboring microbial species are fundamental elements that collectively determine the structure and function of the plant microbiota. However, the molecular basis of such interactions is poorly characterized. Here, we monocolonized *Arabidopsis* leaves with nine plant-associated bacteria from all major phyla of the plant microbiota and profiled co-transcriptomes of plants and bacteria. We detected both common and distinct co-transcriptome signatures among plant-commensal pairs. *In planta* responses of commensals were similar to those of a disarmed pathogen characterized by the suppression of genes involved in general metabolism in contrast to a virulent pathogen. We identified genes that are enriched in the genome of plant-associated bacteria and induced *in planta*, which may be instrumental for bacterial adaptation to the host environment and niche separation. This study provides insights into how plants discriminate among bacterial strains and lays the foundation for in-depth mechanistic dissection of plant-microbiota interactions.

## Introduction

In nature, plants assemble bacterial communities with well-defined taxonomic structures (the plant microbiota) (*1*), which can be harnessed for plant health and survival (*2–4*). How plants discriminate among various bacterial strains and establish strain-specific associations in a community context remain an open question in basic plant microbiota research and is key to facilitating the application of microbiota-based strategies to improve plant health in agricultural settings. Answering this question requires a comprehensive and unified understanding of plant and bacterial responses during their interactions.

Plant responses to microorganisms are controlled by the plant innate immune system, which contributes to the assembly and maintenance of healthy bacterial communities (*5, 6*). A crucial part of the plant immune system is the perception of environmental microbes using cell surface receptors that detect conserved microbial epitopes, termed microbe-associated molecular patterns (MAMPs) (*7*). Recognition of MAMPs triggers defense responses collectively called pattern-triggered immunity (PTI), which can inhibit pathogen growth (*8*). MAMPs such as the bacterial flagellin peptide flg22 are widely conserved in non-pathogenic bacteria as well as pathogenic bacteria (*9*), and some non-pathogenic Proteobacteria strains were shown to trigger defense responses in plant leaves likely via PTI pathways (*10*). On the other hand, diverse microbiota members can suppress PTI triggered by flg22 in roots (*11–13*), which can facilitate colonization by the root microbiota (*11, 13*). Thus, PTI activation by divergent MAMPs and subsequent PTI modulation by plant-associated bacteria might steer plant responses in a bacterial strain-specific manner, contributing to microbiota assembly in plants. A recent study identified, in the aboveground part of plants, a core set of genes induced by phylogenetically diverse endogenous bacteria; some of these genes contribute to plant defense against pathogens (*14*). Therefore, studying common and specific plant responses to diverse bacteria is crucial for our understanding of the role of the plant immune system in the face of both pathogenic and non-pathogenic microbes.

When colonized densely and heterogeneously by various bacterial species, plants might not be able to tailor their responses to individual bacterial strains. Yet, it might be possible that different plant-associated bacterial species respond differently to the same microenvironments created by plants. If so, analyzing plant responses alone does not wholly explain bacterial responses during interactions with hosts. The explanation requires directly interrogating bacterial responses *in planta* at the genome-wide scale. *In planta* bacterial omics approaches, such as transcriptomics, are powerful in understanding bacterial gene functions in the plant microbiome and how plants influence bacterial activities (*15*). To date, however, there are few available *in planta* bacterial transcriptome studies, which focus on pathogenic Proteobacteria strains (*16–21*). It is, therefore, unknown whether plant-associated bacteria have any common or phylum-specific gene expression signatures and what kind of functions are important for their non-pathogenic and sometimes beneficial traits in plants. Integrated analysis of plant and bacterial transcriptome responses is key for building hypotheses about the molecular dialogue between plants and microbiota members.

Here, in monoassociation conditions, we co-profiled the transcriptomes of the model plant *Arabidopsis thaliana* and various bacterial strains isolated from healthy (asymptomatic) plants in nature (hereafter commensal strains), representing all major phyla of the plant microbiota residing in leaves. Commensal strains commonly induced PTI responses in plants, but these differed in intensity. We found examples of both common and strain-specific regulation of commensal gene expression in plants. Bacterial genes enriched in plant-associated strains tended to be induced *in planta*. These included genes involved in sulfur, nitrogen, and carbon transport and metabolism, which were induced *in planta* in a strain-specific manner. This suggests that nutrient status differs for different strains in plants, which may affect bacterial fitness and niche separation. We also observed that plants could elicit different transcriptional responses from different bacterial strains without tailoring their own transcriptional reprogramming. This study provides a framework for dissecting plant-microbiota interactions at the strain level using co-transcriptomics and unravels diverse modes of interactions between plants and commensal bacteria.

## Results

### Co-transcriptome analysis of plants and plant microbiota members

We developed a pipeline to simultaneously investigate host and microbial transcriptomes during plant colonization with a single bacterial strain. We monocolonized *A. thaliana* wild-type Col-0 leaves with individual commensal strains and profiled transcriptomes of plants and bacteria by RNA-seq **(Fig. 1A)**. For *in planta* bacterial RNA-seq, we used a previously developed method with some modifications (Methods). Briefly, bacterial cells are isolated from plant leaves before extracting RNA, followed by rRNA depletion and RNA-seq (*16*). For plant and bacterial RNA-seq, respectively, 18 and nine commensal strains covering all major phyla of the plant microbiota were selected **(Fig. 1B and Table 1)**. Three biological replicates from independent experiments were taken for each condition. We used the same strain IDs as in the original study where these bacterial strains were isolated from wild *A. thaliana* plants (leaves and roots) or soil (*22*). A strain ID indicates the original compartment from which the strain was isolated, but many root/soil isolates can also colonize the shoot, indicating extensive niche overlap (*22*).

**Fig. 1:**
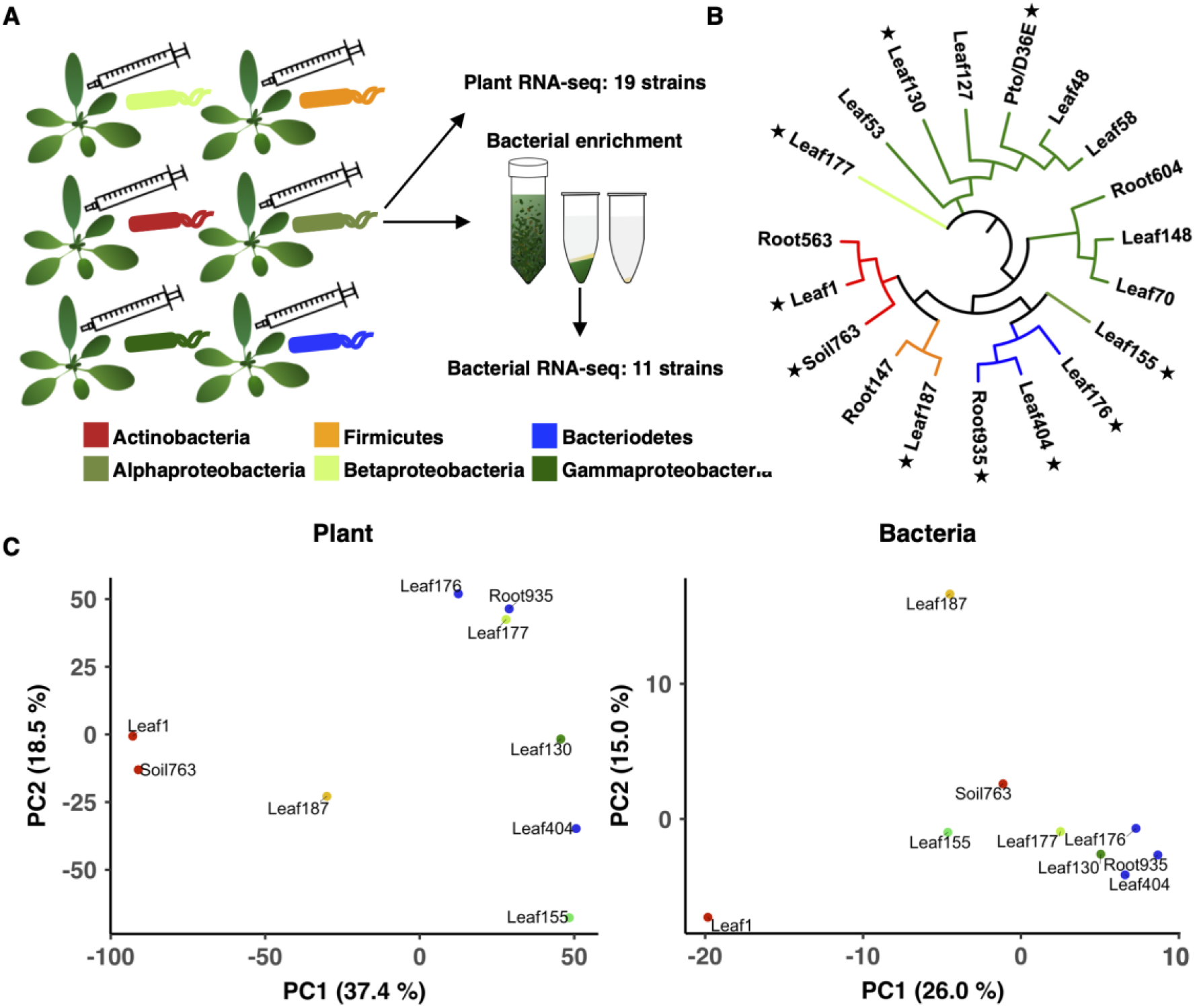
Co-transcriptomics of plants and bacteria. **(A)** Experimental scheme. Individual bacterial strains were syringe-infiltrated into leaves of *A. thaliana* at OD_600_ of 0.5. Leaves were sampled at 6 h post-inoculation. Total RNA was extracted for plant RNA-seq. For bacterial RNA-seq, bacterial cells were isolated from plant leaves before extracting RNA using a method previously reported (*16*). **(B)** Bacterial strains used in this study. Stars indicate the strains used for co- transcriptome analysis. Detailed taxonomic information is shown in Table 1. **(C)** Principal component analysis of gene expression fold changes (FCs) of plants (left: bacteria-inoculated vs. water-inoculated) and bacteria (right: *in planta* vs. *in vitro*). Orthologous groups (OGs) of bacterial genes shared among all strains are used for the analysis. The taxonomic affiliation (phylum/class level) of each strain is indicated with different colors.

**Table 1:** List of bacterial strains used in this study. RNA-seq data (P, plant; B, bacteria) obtained in this study are indicated.

| ID | Phylum | Class | Order | Family | Genus | RNA-seq |
| --- | --- | --- | --- | --- | --- | --- |
| Leaf1 | Actinobacteria | Actinobacteria | Actinomycetales | Microbacteriaceae | Microbacterium | P, B |
| Root563 | Actinobacteria | Actinobacteria | Actinomycetales | Intrasporangiaceae | Janibacter | P |
| Soil763 | Actinobacteria | Actinobacteria | Actinomycetales | Micrococcaceae | Arthrobacter | P, B |
| Leaf176 | Bacteroidetes | Sphingobacteriia | Sphingobacteriales | Sphingobacteriaceae | Pedobacter | P, B |
| Leaf404 | Bacteroidetes | Flavobacteriia | Flavobacteriales | Flavobacteriaceae | Chryseobacterium | P, B |
| Root935 | Bacteroidetes | Flavobacteriia | Flavobacteriales | Flavobacteriaceae | Flavobacterium | P, B |
| Leaf187 | Firmicutes | Bacilli | Bacillales | Exiguobacterium | Exiguobacterium | P, B |
| Root147 | Firmicutes | Bacilli | Bacillales | Bacillaceae |  | P |
| Leaf155 | Proteobacteria | Alphaproteobacteria | Rhizobiales | Rhizobiaceae | Agrobacterium | P, B |
| Leaf177 | Proteobacteria | Betaproteobacteria | Burkholderiales | Burkholderiaceae | Burkholderia | P, B |
| Leaf53 | Proteobacteria | Gammaproteobacteria | Enterobacteriales | Enterobacteriaceae | Erwinia | P |
| Leaf70 | Proteobacteria | Gammaproteobacteria | Xanthomonadales | Xanthomonadaceae | Xanthomonas | P |
| Leaf148 | Proteobacteria | Gammaproteobacteria | Xanthomonadales | Xanthomonadaceae | Xanthomonas | P |
| Root604 | Proteobacteria | Gammaproteobacteria | Xanthomonadales | Xanthomonadaceae |  | P |
| Leaf48 | Proteobacteria | Gammaproteobacteria | Pseudomonadales | Pseudomonadaceae | Pseudomonas | P |
| Leaf58 | Proteobacteria | Gammaproteobacteria | Pseudomonadales | Pseudomonadaceae | Pseudomonas | P |
| Leaf127 | Proteobacteria | Gammaproteobacteria | Pseudomonadales | Pseudomonadaceae | Pseudomonas | P |
| Leaf130 | Proteobacteria | Gammaproteobacteria | Pseudomonadales | Moraxellaceae | Acinetobacter | P, B |
| Pto | Proteobacteria | Gammaproteobacteria | Pseudomonadales | Pseudomonadaceae | Pseudomonas | P, B |
| D36E | Proteobacteria | Gammaproteobacteria | Pseudomonadales | Pseudomonadaceae | Pseudomonas | P, B |

Nine commensal strains, the virulent pathogen *Pseudomonas syringae* pv*. tomato* DC3000 (*Pto*), and its avirulent mutant D36E (36 type III effectors are depleted) were used for co-transcriptome analysis **(Fig. 1B)**. These strains could colonize in the leaf endosphere to various degrees when inoculated on the leaf surface **(Fig. S1)**. To avoid different bacterial population densities to influence plant and bacterial transcriptomes, we syringe-infiltrated bacteria at a defined dose and harvested samples at 6 hours post inoculation (hpi) where the population density of fast-growing *Pto* remained unchanged (*16*).

For plants, we compared gene expression changes between bacteria-inoculated plants and water-inoculated plants **(Fig. 1C; Fig. S2B, left)**. For bacteria, we compared expression changes between *in planta* and *in vitro* (rich media) conditions **(Fig. 1C; Fig. S2A; Fig. S2B, right)**. To directly compare bacterial gene expression patterns among phylogenetically diverse bacterial strains, genes of different strains were grouped based on sequence homology, resulting in 6,823 orthologous groups (OGs) **(Fig. S2E)**. Of these OGs, 454 OGs were shared among all strains **(Data S1)**, indicating that the commensal strains used in this study possess highly diverse gene sets.

Principal component analysis revealed marked differences in both microbial and host transcriptional outputs between plant-commensal pairs, indicating strain-specific interactions between plants and bacteria **(Fig. 1C)**. Interestingly, patterns of transcriptional variation of plants and bacteria were incongruent **(Fig. 1C)**. For instance, we observed similarity between the plant transcriptome changes elicited by different Actinobacteria strains (Leaf1 and Soil763), but these Actinobacteria strains responded highly differently *in planta* **(Fig. 1C)**. Also, *Bacteroidetes* strains (Leaf176, Leaf404, and Root935) showed similar transcriptional changes in plants, but plant transcriptome changes triggered by these strains were distinct **(Fig. 1C)**. These results indicate that plant responses do not fully predict commensal responses and vice versa, pointing to the necessity of co-transcriptome analysis to understand their interactions.

Next, we included pathogens (*Pto* and D36E) in our analysis. The virulent pathogen *Pto* triggered a highly different transcriptome response in plants compared with commensals, whereas plant response to the disarmed pathogen D36E fell between *Pto* and the commensals **(Fig. S2C**), indicating that commensal and disarmed pathogenic bacteria trigger common plant immune responses. Similarly, the gene expression pattern of D36E *in planta* was similar to those of commensals, while the virulent *Pto* showed a distinct pattern (*Pto in planta* resembled commensals *in vitro*) **(Fig. S2D)**. Taken together, co-transcriptome data captured differences in bacterial lifestyles (i.e., virulent vs non-virulent) and revealed commonalities between commensals and a disarmed pathogen.

### Conserved and strain-specific regulation of commensal functions *in planta*

We sought to further analyze bacterial transcriptome data to understand different modes of interactions between different commensal strains and plants. However, the high variability in bacterial genomes complicates a gene-level comparison of bacterial responses among phylogenetically diverse strains **(Fig. S2F)**. One way to overcome this problem is to compare the regulation of bacterial functions rather than of individual genes. Thus, for each strain, we performed functional enrichment analysis on genes significantly up- or down-regulated *in planta* compared with *in vitro* using KEGG functional categories assigned to individual OGs (Methods). Then, enrichment scores (*p* values) for individual KEGG functional categories were summarized for all the strains **(Fig. 2A and Fig. S3A-C)**. We used gene expression fold changes in most analyses to avoid baseline transcriptome differences among strains to confound our analysis (see Discussion for details). Data from *Pto* grown in a minimal medium were included to determine the effect of nutrient availability on gene expression changes. A clear pattern distinguishing virulent and avirulent strains was seen in the process “ribosome” **(Fig. 2A and 2B)**. Genes encoding ribosomal subunits were significantly suppressed *in planta* in all the commensal strains tested and the avirulent pathogen D36E, while these genes were induced in the virulent pathogen *Pto* **(Fig. 2A and 2B)**. This process was also suppressed in *Pto* grown in a minimal medium compared with a rich medium **(Fig. 2A and 2B)**. Since the population density of *Pto* remains unchanged at this time point (*16*), suppression of ribosome-related genes is not the consequence of bacterial growth *in planta*, while these changes could influence on bacterial growth at later time points. Similarly, genes encoding proton ATPases, which are involved in energy production (and possibly alteration of extracellular pH), were induced in *Pto in planta*, but suppressed or not altered in the commensal strains and D36E **(Fig. 2A and 2B)**. Together, these results suggest that commensal strains as well as the disarmed pathogen D36E are metabolically less active *in planta* at an early stage of interactions compared with a virulent pathogen. Since D36E is a mutant of *Pto* lacking PTI-suppressing effector molecules, PTI is likely responsible for suppressing bacterial metabolism *in planta*, and only pathogens can overcome PTI to be metabolically active. Catalase genes were commonly induced in most commensals and D36E at varying degrees but not in *Pto* **(Fig. 2C)**, suggesting that commensals are responding to plant ROS burst, a characteristic PTI response.

**Fig. 2:**
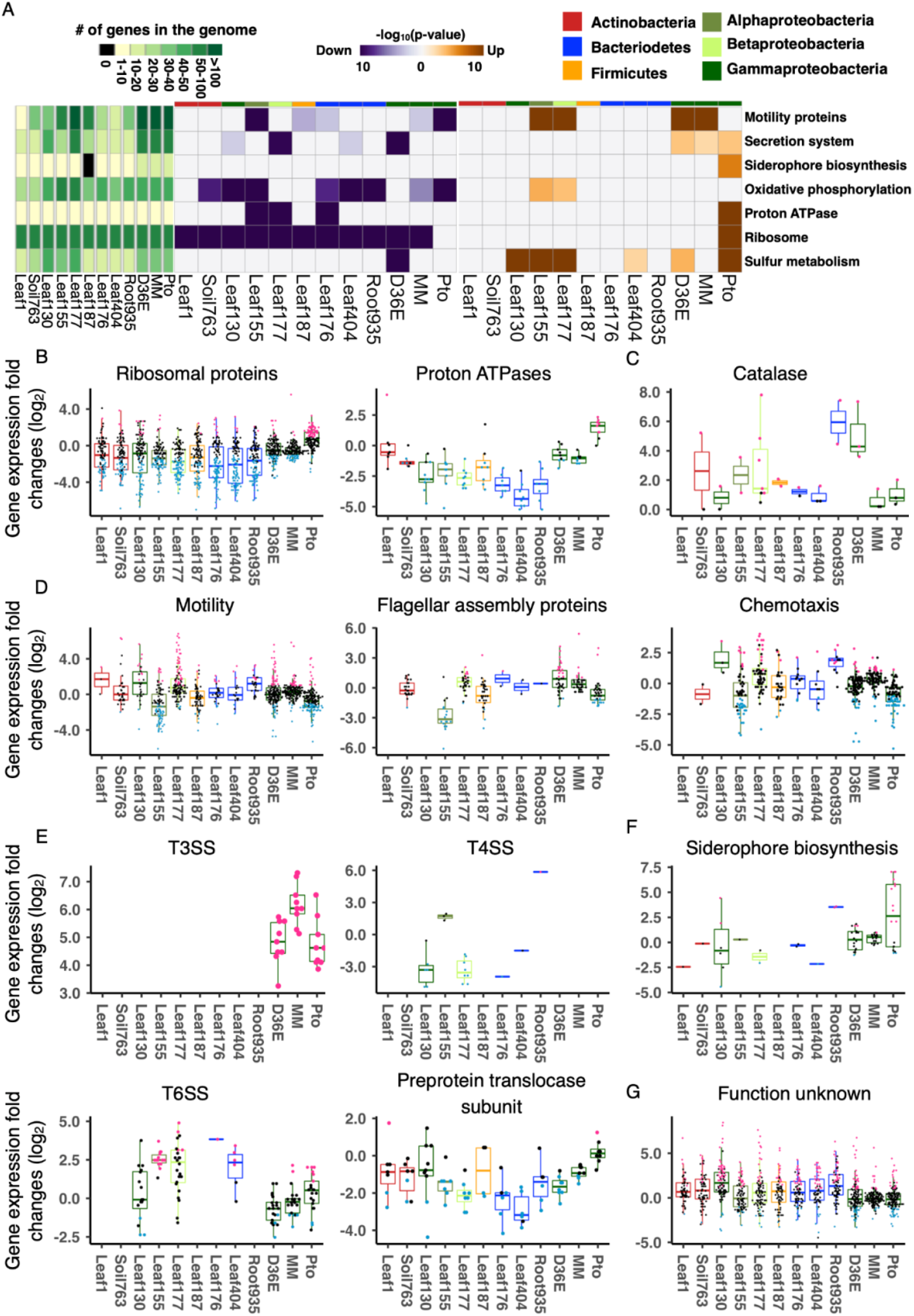
Conserved and strain-specific regulation of bacterial functions in plants. **(A)** KEGG orthology terms enriched in genes that are significantly up- (orange) or down (purple)-regulated *in planta* compared with *in vitro* (rich media). The heatmaps indicate -log_10_ p-value (FDR corrected by Benjamini–Hochberg method). A KEGG orthology can be both significantly up and down-regulated in the same strain. The left green panel shows the number of genes involved in each KEGG orthology term. The top color bars indicate the taxonomic affiliation (phylum/class level) of each strain. See **Fig. S3A** for a more comprehensive list of KEGG orthology. **(B-G)** Expression fold changes (*in planta* vs. *in vitro*) of genes associated with KEGG orthology terms related to (B) general metabolism, **(C)** catalase metabolic pathway, **(D)** motility, **(E)** secretion systems, **(F)** siderophore biosynthesis, and **(G)** unknown functions. T3SS, type III secretion system. T4SS, type IV secretion system. T6SS, type VI secretion system. MM, *Pto* grown in a minimal medium. Results are shown as box plots with boxes displaying the 25th– 75th percentiles, the centerline indicating the median, whiskers extending to the minimum, and maximum values no further than 1.5 inter-quartile range. All individual data points (genes) are overlaid with colors for DEGs (red: upregulated, blue: downregulated, black: non-DEG).

Genes involved in bacterial motility were differentially regulated among bacteria in plants. Many of these genes were suppressed in *Pto in planta* but induced in D36E **(Fig. 2A and 2D)**. Leaf177, a *Burkholderia* (Betaproteobacteria) strain, showed a similar pattern to D36E **(Fig. 2A and 2D)**. However, the *Rhizobiales* (Alphaproteobacteria) Leaf155 more closely resembled virulent *Pto* – a majority of the genes were suppressed *in planta* **(Fig. 2D)**. Motility-related genes can be classified into two major functional categories, flagellar assembly, and chemotaxis. Genes encoding flagellar assembly proteins were globally suppressed *in planta* in Leaf155 as in *Pto*, and many Leaf177 chemotaxis-related genes were induced *in planta* in contrast to *Pto* **(Fig. 2D)**. Thus, physiological processes were differentially regulated among different plant-associated commensal bacteria strains, with some species even exhibiting similarity to a virulent pathogen.

The type III secretion system, an essential component of the virulence of bacterial pathogens, including *Pto*(*23*), was strongly induced in *Pto*and D36E, while these genes were absent in the commensals **(Fig. 2E)**. The type IV secretion system is involved in multiple processes such as translocating proteins and DNA into other cells and bacterial motility (*24*). This process was globally suppressed in Leaf130 and Leaf177, but not in Leaf155 **(Fig. 2E)**. The type VI secretion system is an injection machine involved in bacteria-host and bacteria-bacteria interactions (*25*). This machinery was globally induced in Leaf155 (Agrobacterium) and Leaf404 and partially induced in Leaf177 and *Pto* **(Fig. 2E)**. Lastly, preprotein translocase subunits, which are involved in the bacterial general secretory pathway (*26*), tended to be suppressed in all commensals and D36E, but not in *Pto* **(Fig. 2E)**. Strain-specific regulation of secretion pathways demonstrated here may explain how different strains interact with plant hosts and surrounding microbes. The results showing diverse transcriptional outputs in conserved genes also indicate that presence/absence information of bacterial genes is not sufficient to infer bacterial functions, and *in planta* bacterial gene expression analysis is necessary. It has been shown that genes encoding iron-chelating siderophores are strongly induced in *Pto* upon plant infection, and the induction of these genes is blocked by plant immunity to suppress bacterial growth (*16*) **(Fig. 2F)**. Most of commensal genes associated with KEGG orthology terms related to siderophore biosynthesis were not induced *in planta* resembling D36E, although it is possible that there are other non-characterized genes involved in siderophore biosynthesis in commensals **(Fig. 2F)**. Notably, many genes (3.9-5.1% of the KEGG annotated genes) annotated as “Function unknown” were significantly induced *in planta* in various commensals **(Fig. 2G)**. These functionally unannotated genes induced *in planta* may have unique roles in plant-bacterial interactions.

### Phylum and strain-specific gene expression

To compare expression of individual genes between different strains, we conducted comparative transcriptome analysis focusing on specific phyla **(Fig. S4A and S4C)**. This approach allows more comprehensive comparative transcriptome analysis as more genes are shared among strains within the same phylum. We focused on Bacteroidetes and Proteobacteria, in which 1,422 and 1,122 OGs were shared, respectively (compared to the 454 OGs shared among the nine commensals) **(Fig. S2C, S4A, and S4C)**. Overall, many genes were differentially regulated in a single strain **(Fig. S2C, S4A, and S4C)**. In both of these phyla, a larger number of genes were commonly suppressed among the three strains in plants than commonly induced **(Fig. S4B, S4D, and S5)**. Clusters of genes commonly suppressed *in planta* (clusters 7 & 8 in Bacteroidetes and clusters 1 & 4 in Proteobacteria) were enriched with “ribosome”-related genes **(Fig. S4E)**. “Transporters” were enriched in multiple clusters with various expression patterns **(Fig. S4F and S6A)**, suggesting that transporters can be separated into sub-groups based on regulation in plants. Also, genes annotated as part of a “two-component system” showed strain-specific expression patterns **(Fig. S4G and S6B)**. Taken together, our intraphylum analysis reveals that even relatively closely related commensal strains respond differently *in planta* at the transcriptional level.

### *In planta* bacterial transcriptomics illuminates bacterial adaptation to the leaf environment

Various bacterial functions were differentially regulated in plants in a strain-specific manner. An important question is whether such functional regulation is relevant for bacterial fitness in plants. Comparative genomics is one way to infer bacterial functions associated with adaptation to the plant environment. A previous study compared the genomes of nearly 4,000 plant-associated and non-plant-associated bacterial strains and defined “plant-associated (PA) genes” that are significantly enriched in plant-associated strains (*27*). We analyzed how PA genes are regulated in plants in our transcriptome data. When analyzing the genes shared among nine commensal strains, we observed that genes induced *in planta* tended to be enriched with PA genes, whereas genes suppressed *in planta* tended to be enriched with nonPA genes **(Fig. 3A)**. Remarkably, PA and nonPA genes were significantly enriched with plant-induced and plant-suppressed genes, respectively, for all the commensals, except for the *Firmicutes* strain Leaf187 **(Fig. 3B)**. Therefore, our data suggest that bacterial genes associated with adaptation to the plant environment are indeed activated during the interaction with plants.

**Fig. 3:**
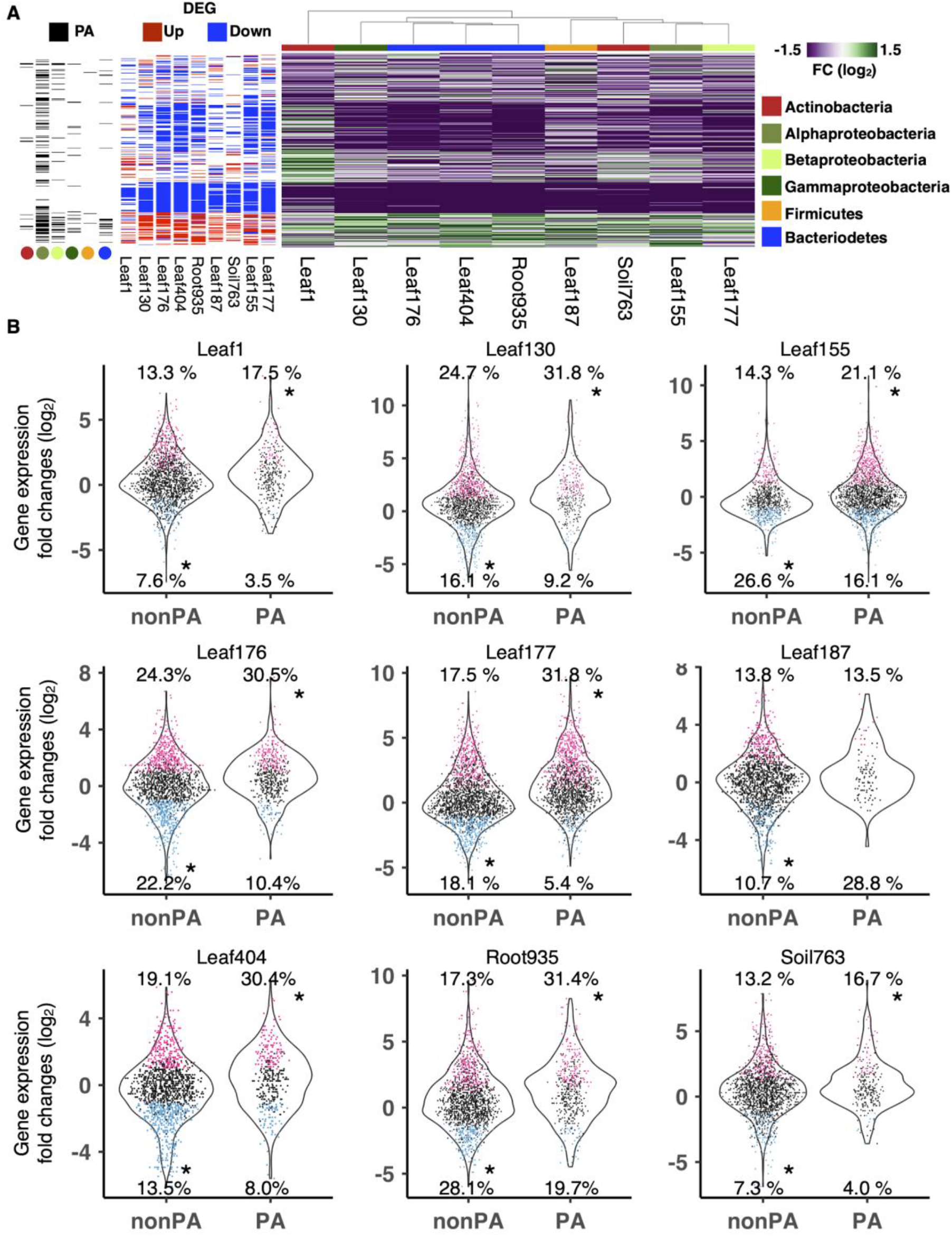
Genes enriched in plant-associated bacteria are induced *in planta*. **(A)** (Right panel) Bacterial gene expression fold changes (FC) in plants compared with *in vitro* (rich media). (Middle panel) Genes differentially expressed *in planta* compared with *in vitro* (|log_2_FC| > 1; FDR < 0.01; two-tailed Student’s t test followed by Storey’s q-value). (Left panel) Genes previously shown to be “plant-associated” (*27*) are shown as black. The bar and dots indicate the taxonomic affiliation (phylum/class level) of each strain. **(B)** Boxplots showing expression changes of plant-associated (PA) and non-plant-associated (nonPA) genes between *in planta* and *in vitro*. Each dot represents a gene. Genes significantly up- or downregulated *in planta* are colored in red and blue, respectively. Asterisks indicate statistically significant enrichment (FDR < 0.05; Hypergeometric test corrected by Benjamini-Hochberg method) of up or down-regulated genes in the PA or nonPA category. The proportion of genes up- or downregulated are shown. For the full expression data with the orthologous group, KEGG annotation, DEG, and PA information, see **Data S3**.

We then performed KEGG functional category enrichment analysis for PA genes induced in plants and nonPA genes suppressed in plants. Ribosome-related genes were conserved among all strains (and are thus nonPA genes) and were generally suppressed in plants **(Fig. 4A)**, which may be a strategy by which plants control bacterial growth. Glycan degradation genes were highly plant-associated and induced in Bacteroidetes strains Leaf176 and Root935 **(Fig. 4A, Fig. S6A)**. Among such genes were homologs of beta-galactosidase, alpha-L-fucosidase, and glucosylceramidase, which can degrade plant cell wall components. Thus, Leaf176 and Root935 may have evolved the ability to degrade the plant cell wall enabling the establishment of favorable niches during plant colonization.

**Fig. 4:**
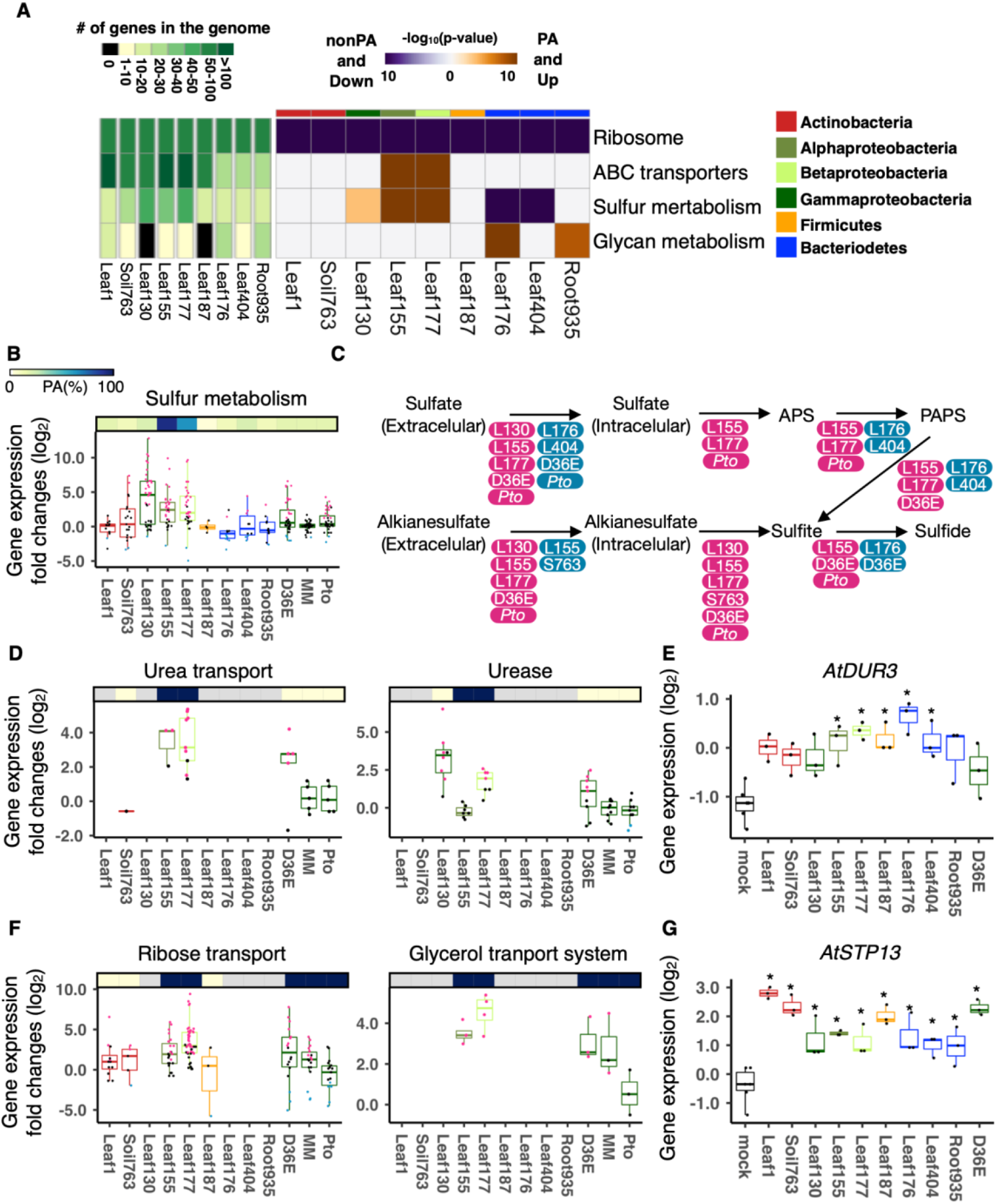
Nutrient acquisition systems are associated with bacterial adaptation to the plant environment in a strain-specific manner. **(A)** KEGG enrichment analysis of genes that are plant-associated (PA) and significantly induced *in planta* compared with *in vitro* (rich media) (orange) and genes that are nonPA and significantly suppressed *in planta* compared with *in vitro* (purple). The left panel shows the number of genes involved in each KEGG orthology term. **(B, D, F)** Expression fold changes (*in planta* vs. *in vitro*) of genes with different functions. The top bars indicate the ratio of PA genes in each strain. All individual data points (genes) are overlaid to the box plots with colors for DEGs (red: upregulated, blue: downregulated, black: non-DEG). **(C)** Suflur metabolic pathways. For each step, a strain name was indicated when the strain has at least one gene significantly induced (red) or suppressed (blue) *in planta*. APS, adenosine 50-phosphosulfate; PAPS, 30-phosphoadenosine-50-phosphosulfate. **(E and G)** Expression of the plant gene **(E)** *AtDUR3* (urea transporter) and **(G)** *AtSTP13* (sugar transporter) based on the RNA-seq data. Asterisks indicate statistically significant difference (|log_2_FC| > 1; FDR < 0.01; two-tailed Student’s t test followed by Storey’s q-value) compared with the mock (water-inoculated) condition. In the box plots, boxes display the 25th–75th percentiles, the centerline indicating the median, whiskers extending to the minimum, and maximum values no further than 1.5 inter-quartile range. For the full expression data with the orthologous group, KEGG annotation, DEG, and PA information, see **Data S3**.

“Sulfur metabolism”-related genes were classified as PA genes and induced *in planta* in the three Proteobacteria strains **(Fig. 4A-4C)**. The sulfur metabolism process includes translocating environmental sulfonate and alkane sulfate into bacterial cells, converting sulfate to APS (adenosine 50-phosphosul-fate), PAPS (30-phosphoadenosine-50-phosphosulfate), sulfite, and then sulfide, which can be converted to amino acids **(Fig. 4C)**. A previous proteomics study showed that the expression of proteins involved in sulfur metabolism and uptake was induced on the leaf surface in two commensal Proteobacteria, *Sphingomonas melonis* and *Methylobacterium extorquens* (*28*). Another study showed the beneficial endophytic bacterium Enterobacter sp. SA187 (Proteobacteria) transcriptionally induces sulfur metabolic pathways *in planta* (*29*). These results suggest that sulfate acquisition is important for the adaptation of commensal Proteobacteria to the plant environment. On the other hand, sulfur metabolism-related genes were not found to be PA genes and were suppressed *in planta* in the Bacteroidetes strains Leaf176 and Leaf404 **(Fig. 4A and 4B)**. Furthermore, the number of genes predicted to be involved in sulfur metabolism was lower in Bacteroidetes strains than in Proteobacteria strains **(Fig. 4A)**. These results may indicate that Bacteroidetes strains are less reliant on sulfur acquisition during plant colonization. As Arabidopsis employs sulfur-containing defense metabolites (*30*), it is also possible that sulfur metabolism-related genes induced in some commensals may serve as a detoxification mechanism rather than a nutrient acquisition mechanism. Genes encoding ABC transporters were PA genes and induced specifically in some Proteobacteria strains *in planta* **(Fig. 4A)**. Among such genes were urea transporters **(Fig. 4D)**. Interestingly, genes encoding ureases, which hydrolyze urea in the bacterial cytoplasm, were also PA and induced in some Proteobacteria strains **(Fig. 4D)**. It has been shown that *Yersinia enterocolitica*, a Gammaproteobacteria strain, can use urea as a nitrogen source (*31*). These results suggest that Proteobacteria (especially Leaf177 and D36E) might use urea as a nitrogen source in the plant apoplast. Genes involved in the nitrate transport system were induced in Leaf130, Leaf155, and D36E *in planta*, but not in *Pto* **(Fig. S7)**, suggesting that some commensal Proteobacteria strains activate nitrogen acquisition systems in plants. Similarly, ribose transporters and glycerol transporters are PA genes and were commonly induced *in planta* in commensal or avirulent Proteobacteria strains (Leaf155, Leaf177, and D36E) but not in the virulent *Pto* **(Fig. 4F)**. Moreover, arabinose and xylose transporters (both monosaccharide transporters) were induced in Leaf177 and D36E *in planta*, but not in *Pto* **(Fig. S7)**. Thus, these Proteobacteria strains may use various types of sugars as carbon sources in plants. The induction of urea and sugar acquisition systems may indicate that commensal bacteria activate nutrient starvation responses in the leaf apoplast. We speculate that plants may sequester nitrogen and carbon sources from the apoplast to limit the growth of commensal and avirulent pathogenic bacteria. In line with this hypothesis, the plant urea transporter *AtDUR3*, which sequestrates urea from the apoplast (*32*), was induced upon inoculation with many commensal strains while suppressed by the virulent *Pto* **(Fig. 4E)**. Since induction of *AtDUR3* has been shown to associate with leaf aging (*32*), it is also possible that these commensals may promote leaf senescence. A previous study showed that plants sequester extracellular sugars by activating the sugar influx transporter *AtSTP13* via the PTI pathway (*33*). Indeed, our plant transcriptome data showed that *AtSTP13* is induced by the commensals as well as D36E and *Pto* **(Fig. 4G)**. On the other hand, *Pto* can induce plant sugar efflux transporters (*34*), which might increase sugar availability in the apoplast and explain why *Pto* did not activate its sugar transporters in plants. A non-mutually exclusive possibility is that the virulent *Pto* switches its metabolic preference to other substrates during successful infection in plants. In summary, we revealed bacterial phylum/strain-specific gene repertoires and gene regulation, which may be actively controlled by plants and drive bacterial niche separation *in planta*.

### Commensals activate plant PTI in a strain-specific manner

As commensal strains showed differing responses in plants, indicating strain specificity in the interactions of plants with bacteria, we further investigated genome-wide plant responses to individual commensals. In addition to the nine commensal strains used for the co-transcriptome analysis, we included nine more commensal strains to enrich our plant transcriptome dataset **(Fig. 1A and Table 1)**. Global gene expression changes (bacteria-inoculated vs. water-inoculated) were qualitatively similar among all commensal strains as well as D36E at 6 hpi **(Fig. 5A)**. Plant gene expression changes triggered by commensals overlapped markedly with responses to flg22 (*35*), a potent PTI inducer. PTI-inducible genes accounted for clusters of genes commonly and strongly induced by most of the commensals (clusters 3/5/7 in **Fig. 5A**). GO enrichment analysis showed that these clusters are enriched with genes related to defense responses (**Fig. 5B**). Thus, commensal strains, when infiltrated into plant leaves, induce common PTI responses.

**Fig. 5:**
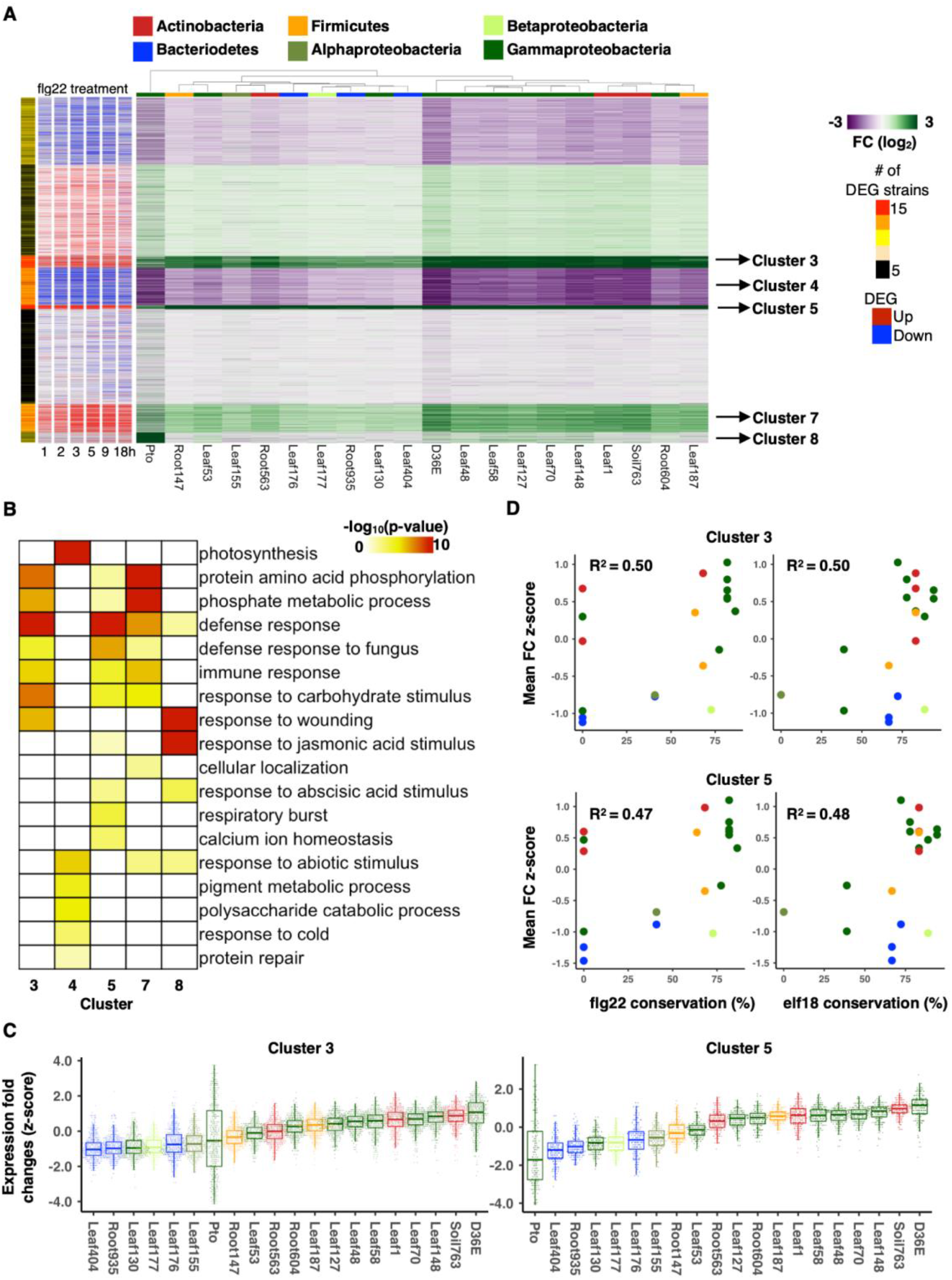
Plant transcriptome responses to phylogenetically diverse commensals. **(A)** (Green/purple heatmap) Gene expression fold changes (FCs) between bacteria-inoculated plants and water-inoculated plants. (Red/blue heatmap) Plant genes significantly induced or suppressed upon flg22 treatment at different time points (*35*). The number of strains causing differential gene expression (|log_2_FC| > 1; FDR < 0.01; two-tailed Student’s t test followed by Storey’s q-value) are indicated in the sidebar (# of DEG strains). DEG, differentially expressed gene. Genes were clustered by k-mean clustering (k = 8). The bars on the heatmaps indicate the taxonomic affiliation (phylum/class level) of each strain. See **Data S4** for gene expression data. **(B)** Gene ontology enrichment analysis for genes in clusters 3, 4, 5, 7, and 8 of **(A)**. -log_10_ p-values (FDR corrected by Benjamini-Hochberg method) were shown. **(C)** Expression fold changes (FC; z-score) of genes in clusters 3 and 5. Results are shown as box plots with boxes displaying the 25th–75th percentiles, the centerline indicating the median, whiskers extending to the minimum, and maximum values no further than 1.5 inter- quartile range. **(D)** Relationships between amino acid (AA) sequence conservation of flg22 or elf18 and normalized expression FCs of genes in clusters 3 and 5. AA sequence conservation of flg22 and elf18 compared with the canonical sequences known to induce strong defense responses in plants (Elf18: SKEKFERTKPHVNVGTIG. Flg22: QRLSTGSRINSAKDDAAGLQIA). The Pearson correlation coefficients are shown. **(C-D)** The same color code was used for the taxonomic affiliation.

The degree of PTI induction varied among strains in a manner that is partly determined by phylogeny: Gammaproteobacteria and Actinobacteria strains induced stronger PTI than Bacteroidetes strains **(Fig. 5C)**. We then investigated the amino acid sequences of the major MAMPs flg22 and elf18 across the different strains. Intriguingly, strains with flg22 and elf18 sequences similar to those known to be particularly potent PTI inducers (*36, 37*) tended to elicit strong PTI induction (gene expression fold changes in clusters 3 or 5) **(Fig. 5D)**. Thus, sequence variation in these MAMPs may partly determine the degree of PTI induced by some of these commensal strains.

### Plant responses are incongruent with bacterial responses in plants

To get deeper insights into the relationships between plant and bacterial gene expression, we measured the correlation between gene expression changes of individual plant genes and shared bacterial OGs using co-transcriptome data of nine commensal strains. To prevent a single outlier strain from impacting correlation scores, we took a bootstrapping approach in which correlations were calculated using all the combinations of eight strains as well as all the nine strains and then combined (Methods) **(Fig. 6A and Fig. S8)**. This analysis revealed that the expression of a majority of plant and bacterial genes is not correlated, further indicating that the plant and bacterial responses are largely uncoupled in our dataset **(Fig. 6B)**. For instance, in many cases, commensal strains that triggered similar plant transcriptional responses (e.g., Soil763 and Leaf1; **Fig. 1C and Fig. 6B**) showed distinct gene expression in plants **(Fig. 1C and Fig. 6B)**. However, a subset of plant and bacterial genes showed a stronger correlation **(Fig. 6B)**. Enrichment analysis of KEGG functional categories showed that expression of bacterial genes annotated as “proton ATPases” and “purine metabolism” positively correlates with plant defense-related genes **(Fig. 6B)**. More specifically, the expression of such bacterial genes was higher when plants showed stronger PTI activation, but a causal relationship between these functions remains elusive. Overall, our data show that plant and bacterial gene expression can be largely uncoupled at an early stage of interaction, indicating that co-transcriptome analysis is required for fully capturing and comparing among various plant-microbe interactions.

**Fig. 6:**
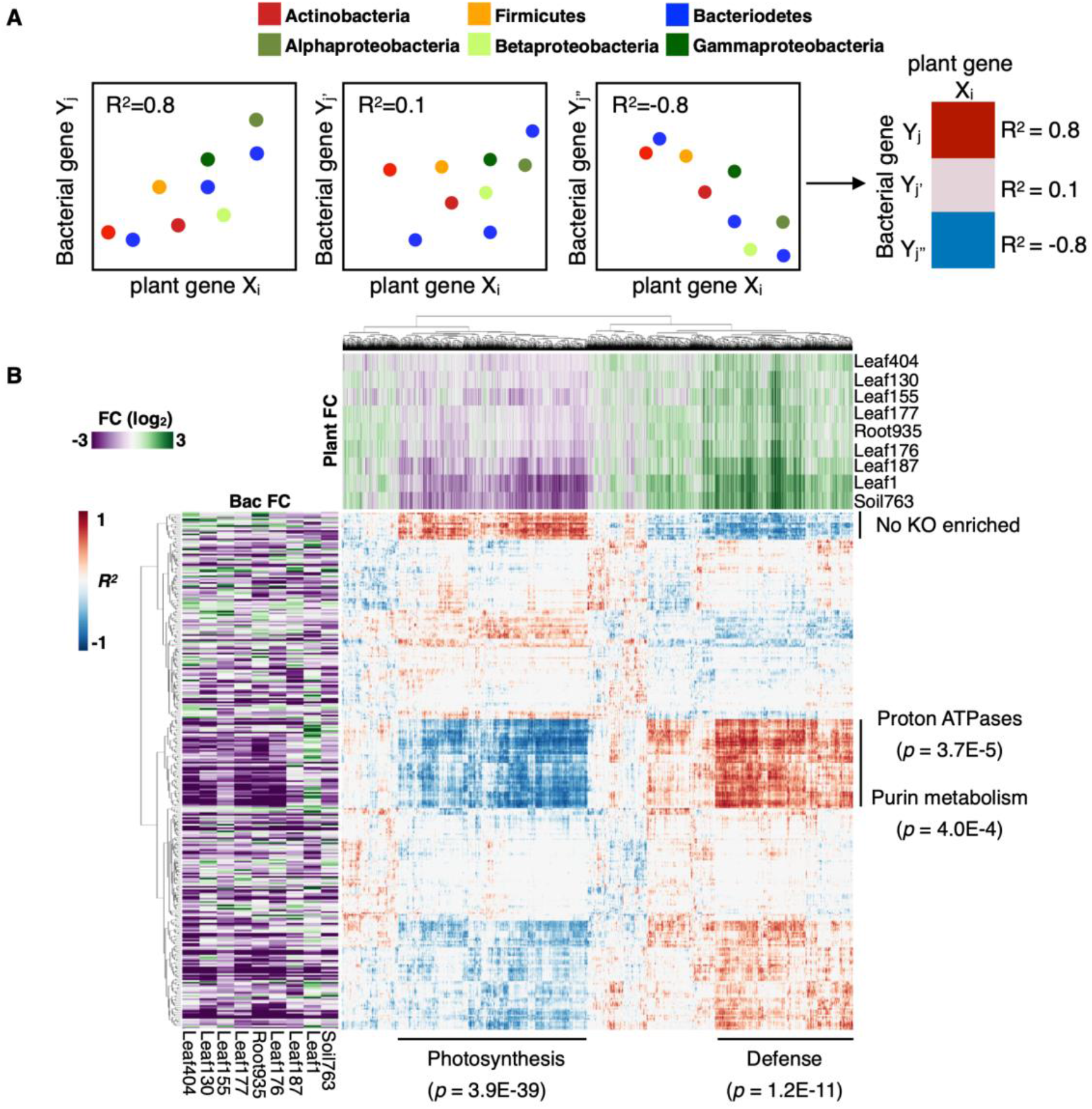
Plant and bacterial transcriptomes are largely uncoupled. **(A)** Schematic diagram of the integration of plant and bacterial RNA-seq data. For each interaction condition, the correlation coefficients between individual plant genes and bacterial OGs were calculated. The correlation coefficient data was corrected by bootstrapping (see Methods and **Fig. S8**) **(B)** A map of correlation coefficients between plant genes and bacterial OGs calculated as described in **(A)**. Rows and columns are bacterial OGs and plant genes, respectively. The top and left heatmaps indicate gene expression FCs of plants and bacteria, respectively. See **Data S5** for the full correlation data. KEGG enrichment analysis was performed for the clusters of plant and bacterial genes with strong correlation.

## Discussion

Previous studies of the plant microbiota have suggested that plants assemble bacterial communities and regulate their functions by interacting with commensals in a strain-specific manner. However, only a limited number of studies have interrogated the responses of plants and commensal bacteria at a genome-wide scale, and thus we do not have a comprehensive understanding of the two-way molecular dialogue between plants and microbiota members. Here, we profiled co-transcriptomes of plants and commensal bacteria in monoassociation using diverse strains covering all major phyla of the plant microbiota. We chose to study an early time point where *in planta* bacterial population density remained unchanged to prevent differential growth across strains that would strongly influence their transcriptomes. Our dataset demonstrated that different commensal strains 1) trigger qualitatively similar yet quantitatively different immune responses in plants and 2) show both common and highly strain-specific responses in plants.

In this study, we primarily analyzed gene expression fold changes (*in planta* vs *in vitro*) to understand bacterial responses during interactions with plants using *in vitro* conditions as a ‘baseline’. Caution is needed when interpreting such data as expression of some genes might be highly different among strains *in vitro*. In our analysis, however, similar patterns were observed even when *in planta* gene expression data alone were used **(Fig. S9)**. Employing multiple baseline conditions in future experiments, such as different media or soil, or performing time course analysis will increase the power of capturing more biologically relevant responses. We syringe-infiltrated bacterial cells into leaves to bypass stomatal entry as different commensals might have different abilities to access the apoplast. Transcriptomes were profiled at 6 hpi, where the population density of even the virulent pathogen *Pto* has not yet increased (*16*), and thus we assumed that the population density of commensals remained the same at this time point. Therefore, our experimental setup allowed us to characterize strain-specific co-transcriptomes under controlled conditions without the influence of stomatal immunity and differences in the sizes of bacterial populations. It is, however, important to note that transcriptome analysis under more natural conditions will reveal additional layers of plant-microbiota interactions. This requires technological innovations that enable *in planta* transcriptome analysis of bacteria with much smaller populations.

We found that suppression of genes related to general metabolic activity and energy production *in planta* is a common trait among phylogenetically diverse commensals, in marked contrast to a virulent pathogen, which elicited the opposite response **(Fig. 2)**. Note that we sampled bacteria at the time point when they did not start multiplying. Thus, higher metabolic activity of the virulent pathogen was not simply due to their active growth. PTI was commonly induced by the commensal strains **(Fig. 5A)**, suggesting that plant immunity might act to keep commensal metabolic activity in check to avoid overgrowth. This notion is in line with a previous finding that commensals can proliferate in an unrestrained manner in the leaf apoplast of plant mutants lacking key immune components (*5, 38*). Further transcriptome analysis of commensals in immunocompromised plants and different environmental conditions will unravel how different immune pathways tailor their responses to effectively control commensal growth and function. Also, testing various other pathogens would be important to reveal lifestyle-dependent transcriptome signatures of bacteria in plants.

We provide evidence that bacterial genes enriched in the genomes of plant-adapted strains are frequently induced *in planta* **(Fig. 3A and 3B)**, suggesting that those genes which enable bacteria to thrive in the plant environment are indeed activated in plants. This finding is somewhat in contrast to a previous study that showed gene expression of a bacterial pathogen *in planta* does not correlate with fitness scores determined by transposon insertion mutagenesis (*39*). Importantly, loss-of-function screening with single mutants has limitations in assigning gene function owing to functional redundancy. In this case, a gain-of-function assay is a complementary, albeit also limited, approach. For instance, *in planta* bacterial transcriptome data could predict bacterial genes that contribute to bacterial growth *in planta* when overexpressed (*16, 40*).

We found that processes involved in the uptake of nutrients, such as sulfur, urea, and sugars, were enriched in plant-associated Alpha- and Betaproteobacteria (L155 and L177) and induced *in planta* **(Fig. 4)**. We also observed that plants induce transporters that could sequester urea and sugars from the apoplast during interactions with commensals **(Fig. 4E and 4G)**, which potentially affects bacterial nutrient acquisition processes and eventually bacterial fitness in plants. Notably, we found many other bacterial nutrient transporters to be regulated *in planta* in a strain-specific manner **(Fig. S7)**. The results imply that different commensals experience distinct nutrient status in the plant apoplast, which might affect bacterial fitness *in planta*. Alternatively, different commensals may respond differently to the same nutrient status. Our co-transcriptome data sets the stage for investigating whether plants control nutrient availability for particular strains to drive bacterial niche separation in plants and shape the plant microbiota.

We did not observe a strong association between gene expression changes in plants and commensals **(Fig. 1C; Fig. 6B)**, implying that similar programs of plant gene expression could divergently affect different commensals. This seems reasonable, given that plants have to deal with complex bacterial communities residing in an area smaller than the plant cell where plant immune responses elicited by a microbe can affect other microbes. In such a situation, recognition of microbes by immune receptors might be insufficient for distinguishing different microbes. Divergent effects of plant immune outputs on bacterial responses may enable plants to selectively host specific strains from complex microbial sources. In the future, time-course analysis of a larger number of strains combined with plant and bacterial genetics will facilitate the prediction of mechanistic links between plant and bacterial responses.

Since plants used in this study were not grown in a strictly sterile condition, we do not exclude the possibility that the pre-existing plant microbiota influenced plant and bacterial responses. However, influence of the pre-existing microbiota on data interpretation was minimized by including the mock control (for plant transcriptomics), randomizing sampling, and taking three independent replicates (for plant and bacterial transcriptomics). We demonstrated that an *in planta* bacterial transcriptome approach can be applied to all major phyla of the plant microbiota, opening a new avenue for *in planta* transcriptome analysis of synthetic communities that are generated by mixing bacterial strains in a desired manner. This *in planta* bacterial metatranscriptome approach together with individual bacterial transcriptomes can capture more complex traits such as microbe-microbe interactions, which are important to understand the functions of the microbiota as a community.

This study provides a wealth of information regarding gene regulation of both plants and commensals during monoassociations. In **Fig. S10-13**, we provided additional insights on the regulation of genes related to diverse functions, including biosynthesis/metabolism of various compounds, transporters, and nucleic acid regulation. Notably, we found many genes with unknown functions to be dynamically regulated in commensals during interactions with plants **(Fig. 2G)**. To explore these commensal functions, it will be critical in the future to link bacterial transcriptome responses to bacterial niche preference and reproductive fitness in plants. Our co-transcriptome dataset will provide a robust platform for hypothesis-driven functional investigation of plant and bacterial genes that play critical roles in plant-microbiota interactions.

## Acknowledgments

We thank Paul Schulze-Lefert and Julia Vorholt for providing the commensal strains, Alan Collmer for providing the Pto DC3000 D36E strain, the Max Planck Genome Centre for sequencing support, Dieter Becker, Ryosuke Kanaoka, Shinpei Shimokawa, and Ying Tang for research assistance, and Neysan Donnelly, You Lu, and Natsuki Ohmae for critical comments on the manuscript. This work was supported by the Fundamental Research Funds for the Central Universities (Program No. 2662020ZKPY009) (to K.T.), Joint Funding of Huazhong Agricultural University and Agricultural Genomics Institute at Shenzhen, Chinese Academy of Agricultural Sciences (SZYJY2021007) (to K.T.), the Huazhong Agricultural University Scientific & Technological Self-innovation Foundation (to K.T.), the Max Planck Society (to R.G.-O. and K.T.), a German Research Foundation grant (SPP2125) (to R.G.-O. and K.T.), a predoctoral fellowship from the Nakajima Foundation (to T.N.), and a Chinese Scholarship Council PhD stipend (CSC Student ID 201808440401) (to Y.C.).

## Author contributions

T.N. and K.T. designed the research. T.N., Y.C., F.E., and Y.T. performed experiments. T.N., E.D., and R.G.O. performed analysis. T.N. and K.T. wrote the paper with input from all authors.

## Data availability

The RNA sequencing data used in this study are deposited in the National Center for Biotechnology Information Gene Expression Omnibus database (accession no. GSE150422). Key data and scripts are available at https://github.com/tnobori/co-transcriptomics.

## Methods

### Plant materials and growth conditions

The *Arabidopsis thaliana* accession Col-0 plants were grown in a chamber at 22°C with a 10-h light period and 60% relative humidity for 24 days and then in another chamber at 22°C with a 12-h light period and 60% relative humidity. For all experiments, 31- to 33-day-old plants were used.

### Bacterial strains

Commensal strains were previously isolated from wild *A. thaliana* plants (*22*) (http://www.at-sphere.com/) **(Table 1)**. The *Pto* mutant D36E was previously described (*41*). Bacterial strains were cultured at 20°C (commensal strains) or 28°C (*Pto* and D36E) at 200 rpm in liquid 50% TSB medium (Sigma-Aldrich, USA).

### Sampling of bacteria *in vitro*

Commensal strains were pre-grown on solid 50% TSB plates for 2-4 days and then grown in liquid 50% TSB medium (starting at OD_600_ = 0.1) and harvested at the late log phase, which was determined by *in vitro* time course growth assays **(Fig. S14)**. 0.1 volume of the stop buffer (95% EtOH, 5% Phenol) was added to bacterial cultures before centrifuging to collect bacterial cells. Target OD_600_ for each strain: Leaf1 = 0.7 (harvested after 6 h), Leaf130 = 1.2 (harvested after 4 h), Leaf155 = 0.5 (harvested after 6 h), Leaf176 = 0.9 (harvested after 8 h), Leaf177 = 0.6 (harvested after 8 h), Leaf187 = 0.8 (harvested after 4 h), Leaf404 = 0.6 (harvested after 4 h), Root935 = 0.8 (harvested after 4 h), Soil763 = 1.8 (harvested after 7 h).

### Bacterial inoculation to plant leaves and sampling

Commensal strains were grown in the liquid 50% TSB medium. For each strain, multiple cultures were prepared with different bacterial densities to ensure that unsaturated cultures were used for experiments. Bacterial cells were harvested by centrifugation, washed twice with sterile water, and resuspended in sterile water to OD_600_ of 0.5. Plants grown in pots were randomized before bacterial inoculation. Leaves were harvested 6 h after inoculation. At this time point, (*16*)did not show increased population density (*16*), thus the population density of slow-growing or non-growing commensals likely remained unchanged. For bacterial RNA-seq, 80–100 *A. thaliana* leaves (four fully expanded leaves per plant) were syringe-inoculated with bacterial suspensions using a needleless syringe. For plant RNA-seq, approximately six leaves (two fully expanded leaves per plant) were treated. Mock control (water infiltration) was included in every plant RNA-seq experiment. Leaves were harvested at 6 hours after inoculation. Sampling took approximately 5 min per genotype. Leaves were immediately frozen in liquid nitrogen and stored at -80°C. Three biological replicates from independent experiments were taken for each condition of plant and bacterial RNA-seq.

### Sequencing library preparation and RNA sequencing

*In planta* bacterial transcriptome analysis was conducted as described previously (*16*) with slight modifications. Briefly, bacteria-infected leaves were coarsely pulverized and released into bacterial isolation buffer (9.5% ethanol, 0.5% phenol, 25 mM TCEP (tris(2-carboxyethyl)phosphine) pH 4.5 adjusted with NaOH) at 4°C, filtered, and centrifuged to isolate bacterial cells from plant cells. The original RNA extraction method based on chemical lysis of bacterial cells by TriFast (*16*) did not work for some bacterial strains, thus we used FastRNA PRO™ BLUE KIT (MP Biomedicals), which involves mechanical cell lysis. rRNA was depleted to enrich mRNA, and the cDNA libraries were prepared using Ovation Complete Prokaryotic RNA-seq kit 1-8 (NuGEN).

For plant RNA-seq, RNA was extracted with FastRNA PRO™ KIT with Lysing Matrix E (MP Biomedicals), and DNA was digested with TURBO DNase (Ambion). RNA quality was determined using a 2100 Bioanalyzer (Agilent Technologies, USA). Initially, 500 ng total RNA was used for polyA enrichment with the Biolabs). Subsequent library preparation was performed with NEBNext® Ultra™ II Directional RNA Library Prep Kit for Illumina® (New England Biolabs) according to the manufacturer’s instructions.

Libraries were immobilized and processed onto a flow cell with cBot (Illumina) and subsequently sequenced on the HiSeq3000 system (Illumina) with 1 x 150 bp single reads. Primary data analysis (incl. image analysis, cluster identification, base calling, assignment of quality scores) has been performed with RTA (real-time analysis software; Illumina) installed on the sequencing platform.

For bacterial and plant samples, approximately 10 and 30 million reads, respectively, were obtained. Bacterial reads were mapped onto the corresponding bacterial genomes (*22*) using Bowtie2 (*42*). Plant reads were mapped onto the *Arabidopsis* genome (TAIR10) using HISAT2 (*43*). Mapped reads were counted with the Python package HTSeq (*44*). The RNA-seq data used in this study are deposited in NCBI Gene Expression Omnibus database (accession no. GSE150422).

### Raw data

Raw RNA-seq count and bacterial gene annotation files are available at https://github.com/tnobori/co-transcriptomics.

### Data analysis – plant RNA-seq

The statistical analysis of the RNA-seq data was performed in the R environment. Genes with average counts < 5 were excluded from the analysis. The count data were TMM-normalized and log-transformed using the function calcNormFactors in the package edgeR (*45*) and the function voomWithQualityWeights in the package limma (*46*), respectively. To each gene, a mixed linear model was fitted by using the function lmFit in the limma package (*46*). Note that mock control (water infiltration) was included in every plant RNA-seq experiment. The eBayes function in the limma package was used for variance shrinkage during the calculation of the p-values. The false discovery rate (FDR; the Storey’s q-values) was calculated using the qvalue function in the qvalue package (*47*). Genes with q-value <0.01 and |log_2_ fold change| > 1 were defined as differentially expressed genes. The prcomp function was used for principal component analysis. Heatmaps were created with the pheatmap function in the R environment. Enriched GO terms were identified using the BiNGO plugin for Cytoscape (*48*). Scatter plots and box plots were generated using the R-package ggplot2.

### Data analysis – bacterial RNA-seq

#### Bacterial phylogenetic analysis

The bacterial genomes were searched for the bacterial small ribosomal subunit 16S rRNA gene using RNAmmer (*49*). Next, a multiple sequence alignment was performed using Clustal Omega (*50*) with default parameters. Finally, we employed FastTree (*51*) to build a maximum-likelihood phylogeny using the gamma time reversible substitution model (GTR) of DNA evolution. This tree was visualized (Fig. 1B) using the interactive Tree of Life (*52*).

#### Orthologous gene prediction and KEGG annotation

*De novo* orthology prediction was performed by using OrthoFinder (*53*) with default parameters on the predicted protein coding sequences extracted from the bacterial genome assemblies. OrthoFinder is a reference-free algorithm that uses pair-wise protein alignments followed by a graph-clustering step to infer orthologous relationships, and has been shown to have a higher accuracy than traditional approaches such as identifying best-bidirectional hits (doi: 10.1093/gbe/evt132). Next, individual genes were annotated with the KEGG database as a reference (*54*) using the blastkoala webserver (Prokaryotes group) (*55*). Subsequently, orthologous genes were assigned a single KEGG orthology annotation by majority vote of individually annotated sequences in each group. The genomes of the commensal strains were previously reported (*22*) and are available at our GitHub repository (https://github.com/tnobori/co-transcriptomics).

#### Data normalization and visualization

RNA-seq data were normalized for each strain. After omitting genes with average count < 5, count data was TMM-normalized and log-transformed as described above. Genes with FDR <0.01 (corrected by Benjamini-Hochberg method) and |log_2_ fold change| > 1 were defined as differentially expressed genes. Commensal genes were annotated with OGs to integrate gene expression data of different strains. When multiple genes are annotated with the same OG, the mean expression value was taken. Data visualization was performed as described above. UpSet plots were generated in the R environment using the package UpSetR (*56*).

### KEGG orthology enrichment analysis (related to **Fig. 2A and Fig. S3A**)

A custom KEGG orthology database was created by taking only functional terms encoded in at least one bacterial genome (downloaded in January 2019). For each strain, a list of KEGG orthologies was generated by subsetting the corresponding KEGG IDs from the custom KEGG orthology database **(Data S6)**. KEGG orthology enrichment test was performed using a hypergeometric test (FDR corrected by Benjamini-Hochberg method). KEGG orthologies with FDR < 0.01 and containing more than three genes were defined as significantly enriched KEGG orthologies. An R script and KEGG orthology databases are available at https://github.com/tnobori/co-transcriptomics.

### Generating plots of genes with various functions (related to Fig. 2B-G, Fig. 4B-D and F, Fig. S3B and C, Fig. S6, Fig. S7, Fig. S9-12)

Bacterial genes were selected by KEGG pathway annotations or keyword searches from KEGG BRITE annotations. R scripts for this analysis are available at https://github.com/tnobori/co-transcriptomics.

### Intersecting plant-associated bacterial genes and differentially regulated genes *in planta* (related to **Fig. 3**)

In a previous study (*27*), comparative genomics analyses defined “plant-associated (PA) genes” for each phylum/class using multiple statistical tests. The study defined two groups of Actinobacteria (Actinobacteria1 and Actinobacteria2). The Actinobacteria strains used in the present study are all Actinobacteria1). We defined genes that passed at least one statistical test as “PA genes” and the others were defined as nonPA genes. An R script and PA-gene datasets for this analysis are available at https://github.com/tnobori/co-transcriptomics.

### MAMP conservation analysis (related to **Fig. 5D**)

Canonical flg22 and elf18 sequences were blasted against the bacterial genomes using blastp (*57*) with standard settings. The results of these homology searches were filtered by retaining hits covering at least 90% of the length of the MAMP sequence in the alignment and subsequently retrieving the alignment with the highest percentage identity.

### Integration of plant and bacterial RNA-seq data (related to **Fig. 6**)

Co-transcriptome fold change data (bacteria: *in planta* vs. *in vitro*; plants: bacteria vs. mock) of nine strains were used for this analysis. Plant genes whose expression was significantly changed by at least one strain were used. Pearson’s correlation coefficients between individual plant genes and bacterial OGs were calculated. The same analysis was performed for all the combinations of eight strains (bootstrapping). Among these 8-strain and 9-strain datasets, the weakest correlation coefficient value was used for each combination of a bacterial OG and a plant gene (Fig. S8). An R script and plant/bacterial gene expression datasets for this analysis are available at https://github.com/tnobori/co-transcriptomics.

### Determination of bacterial colony forming units (related to Fig. S1)

Bacterial colonization of the leaves was determined following a previous study (*5*) with slight modifications. The Cl_2_-gas-sterilized seeds were stratified for 2 days at 4°C, sown on half Murashige & Skoogs (MS, Duchefa-Biochemie, MO255.0050) agar medium with 1% sucrose, and allowed to germinate for 5 days. Seedlings of the same physiological state were transplanted on half MS agar medium and were grown for another 9 days (a total of 2 weeks) prior to inoculation with bacteria. One day before inoculation, bacterial cultures were grown on half TSB for 24 hours at 22°C with 200 rpm shaking. On the day of inoculation, bacterial cells were harvested by centrifugation at 3000 rpm for 15 min, washed twice with sterile water, and then finally suspended in 10 mM MgCl_2_. The resulting bacterial suspensions were diluted to a final OD_600_ of 0.5 with sterile water and with this, each plate of 2-week-old seedlings was flood-inoculated for 1 min, drained, and allowed to dry for 15 min. Plants were then grown for 3 days and 2-3 leaves of the same physiological state were harvested aseptically and weighed. To quantify bacteria in the endophytic compartment, leaves were surface-sterilized with 75% ethanol for 30 seconds and washed twice with sterile water, and the leaves were homogenized on 10 mM MgCl^2^ buffer using TissueLyserll (Qiagen) with the frequency of 30 s^-1^ for 5 min. The samples were then serially diluted (10^0^ to 10^5^) and spread-plated on 0.5x TSB agar medium. Plates were incubated at ambient temperature, colonies were observed and counted for 1-3 d and colony forming units were expressed per mg FW. The total compartment was assayed similarly but without surface sterilization.

Data S1 OG distribution

Data S2 Expression of all OGs

Data S3 List of PA and nonPA KEGG orthologies

Data S4 Plant RNA-seq data

Data S5 Correlation matrix of plant and bacterial transcriptomes

Data S6 KEGG orthology database for each strain

## Supplementary figures

**Fig. S1.**
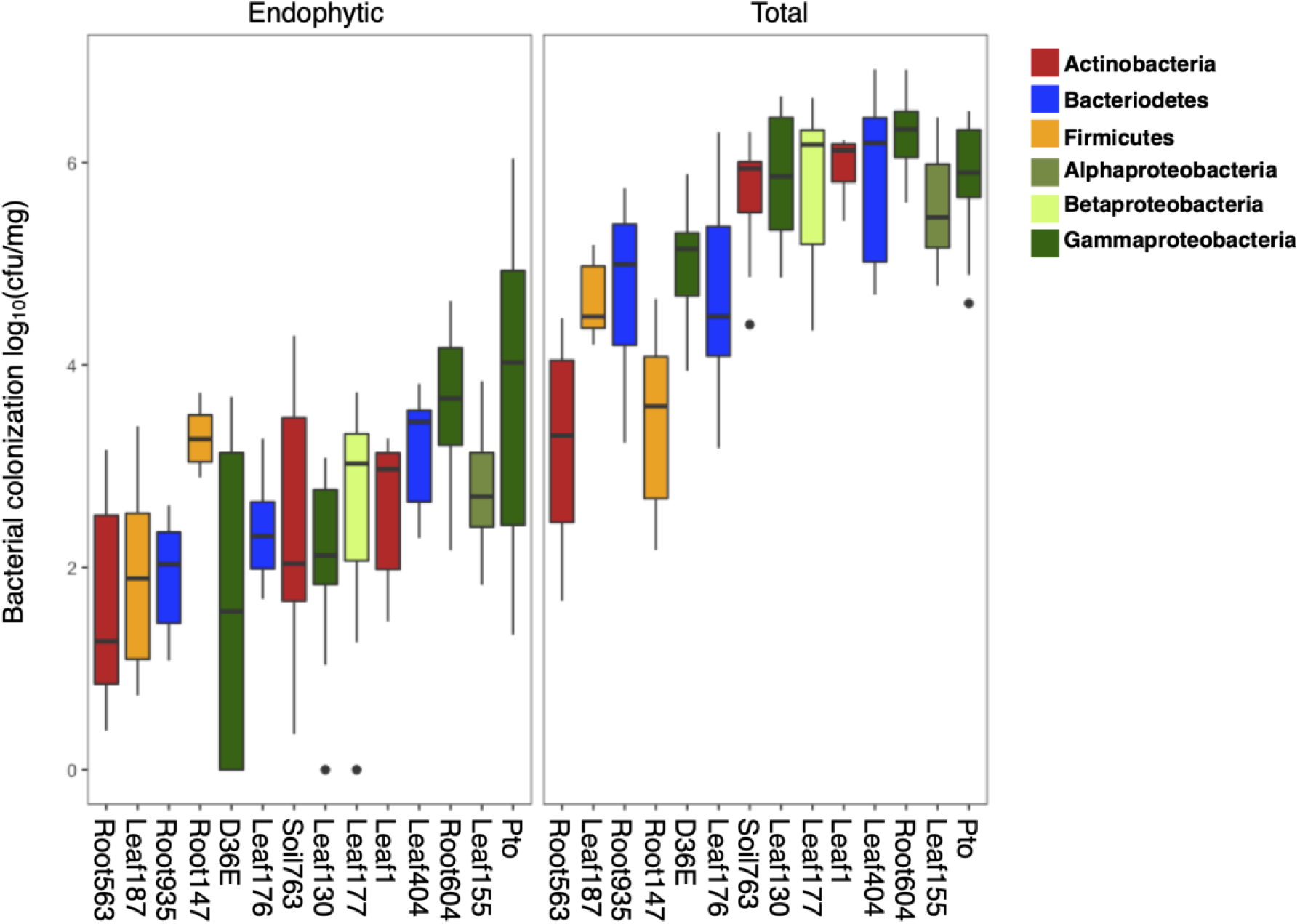
Commensal strains can colonize in leaf endosphere. Bacteria were flood-inoculated to three-week-old *A. thaliana* Col-0 at OD_600_ = 0.5. Bacterial growth was measured three days after inoculation. (Left) Bacteria that entered the leaf endosphere and persisted or grew were counted after washing and sterilizing the leaf surface. (Right) total bacteria (endophytic + epiphytic) were counted without any surface washing and sterilization (see Methods).

**Fig. S2.**
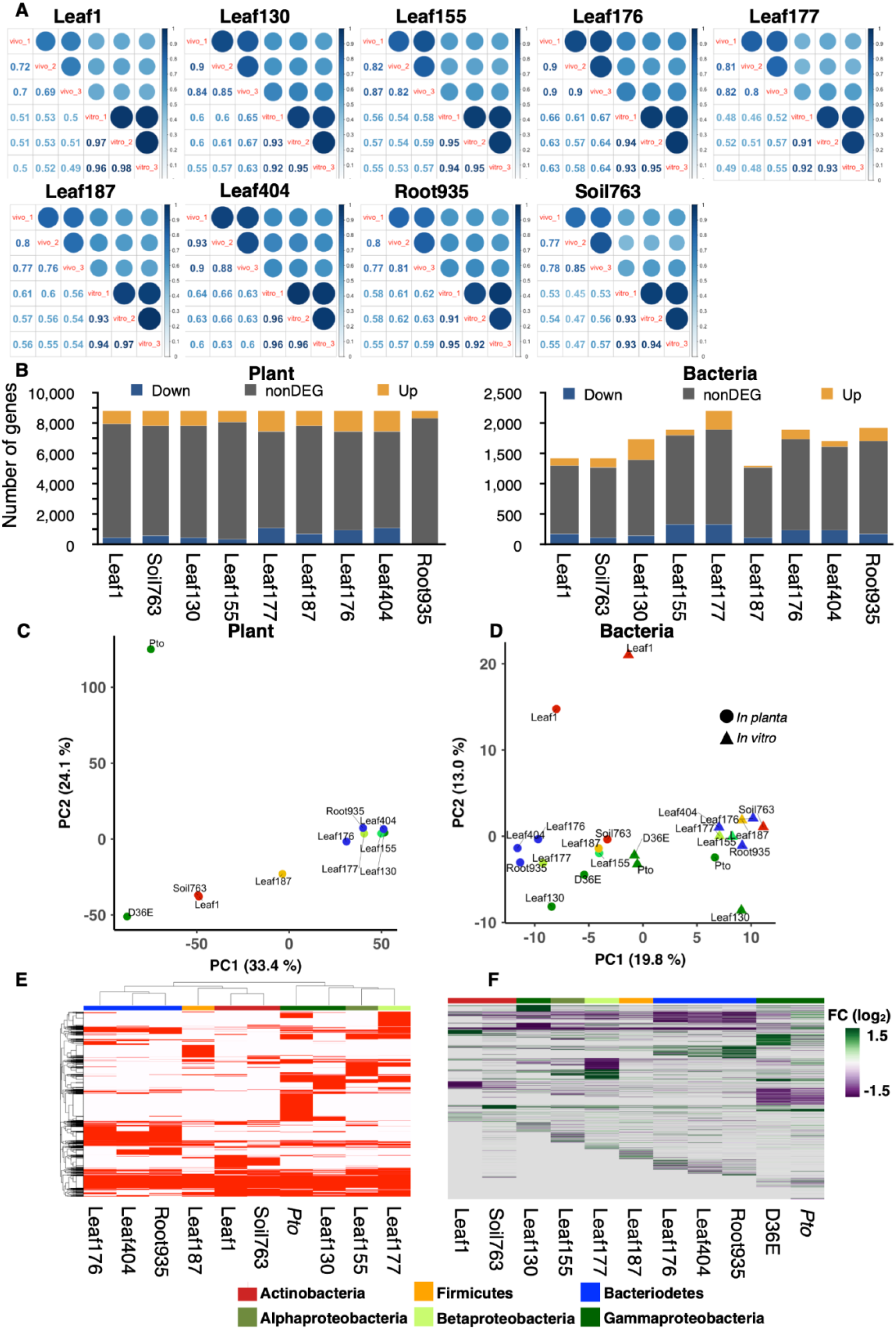
Transcriptome analysis of bacteria. **(A)** The correlation plot of each replicate of bacterial RNA-seq data for individual strains. vivo: bacteria in plants. vitro: bacteria in rich media. **(B)** The number of genes analyzed in plant or bacterial RNA-seq. Differentially regulated genes (|log_2_FC| > 1; FDR < 0.01; two-tailed Student’s t test followed by Storey’s q-value) are colored in blue or yellow. Plant: bacteria-inoculated vs. water-inoculated. Bacteria: *in planta* 6 h vs. *in vitro* (rich media). Bacterial strains used for co-transcriptomics are shown. Plant and bacterial genes with low expression levels across samples were removed. **(C)** Principal component analysis of gene expression fold changes (FCs) of plants inoculated with bacteria used for the co-transcriptome analysis (bacteria-inoculated vs. water-inoculated). **(D)** Principal component analysis of bacterial gene (orthologous group) expression *in planta* and *in vitro*. **(E)** Genes that are present in each strain are colored in red. See **Data S1** for the gene presence-absence table. **(F)** Gene expression fold changes (*in planta* vs. *in vitro*) of bacteria. The data of *Pto* and D36E are from a previous study (*16*). Genes not detected or missing in the genome are shown in gray. See **Data S2** for gene expression data. **(C-F)** The taxonomic affiliation (phylum/class level) of each strain is indicated with different colors.

**Fig. S3:**
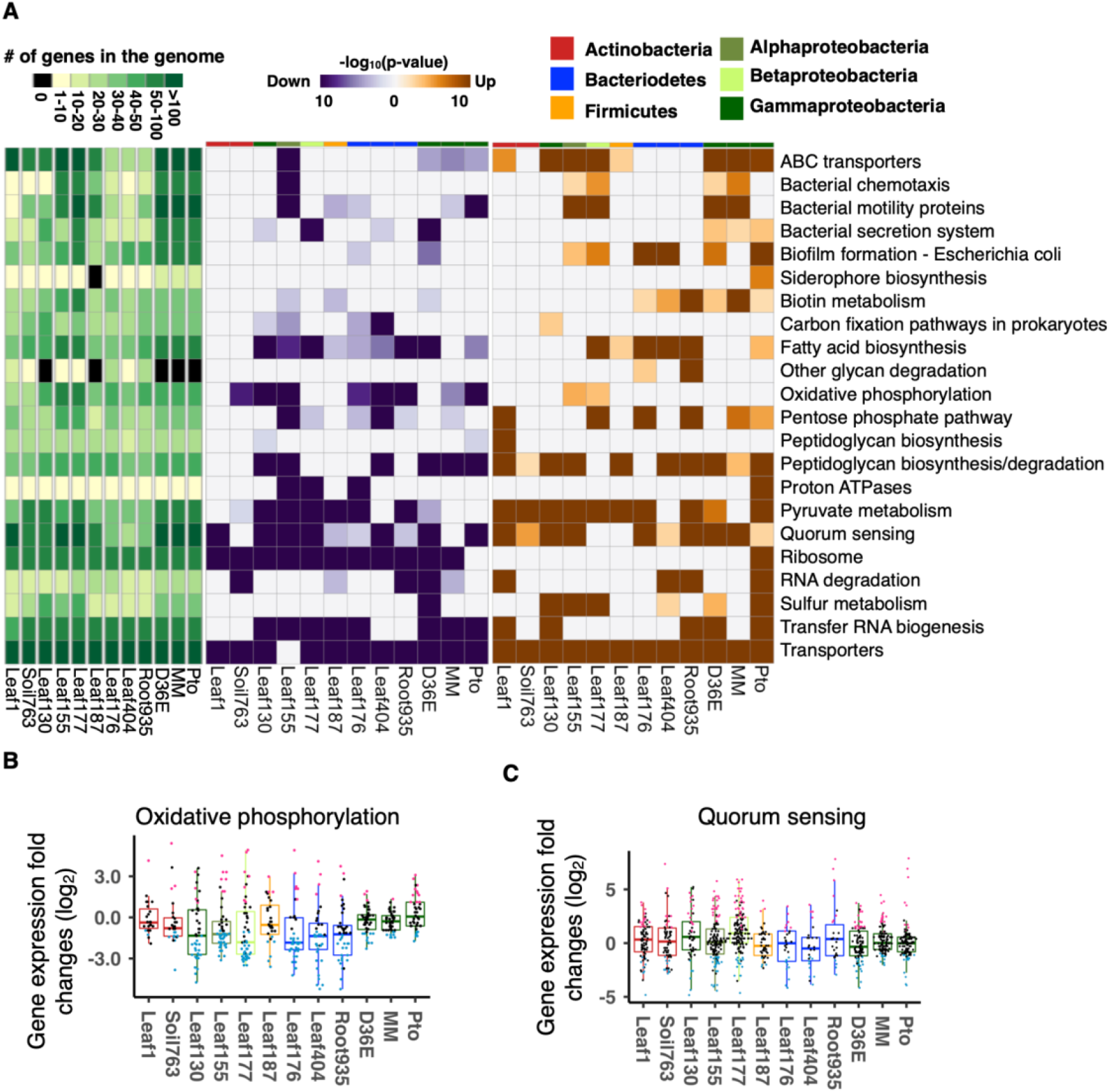
Conserved and strain-specific regulation of bacterial functions in plants. **(A)** KEGG orthology terms enriched in genes that are significantly up- (orange) or down (purple)-regulated *in planta* compared with *in vitro* (rich media). The heatmaps indicate -log_10_ p-values (FDR corrected by Benjamini-Hochberg method). The top color bars indicate the taxonomic affiliation (phylum/class level) of each strain. **(B and C)** Expression fold changes (*in planta* vs. *in vitro*) of genes involved in **(B)** Oxidative phosphorylation and **(C)** Quorum sensing. MM, *Pto* grown in a minimal medium. Results are shown as box plots with boxes displaying the 25th–75th percentiles, the centerline indicating the median, whiskers extending to the minimum, and maximum values no further than 1.5 inter-quartile range. Box color indicates the taxonomic affiliation (phylum/class level) of each strain. All individual data points (genes) are overlaid with colors for DEGs (red: upregulated, blue: downregulated, black: non-DEG).

**Fig. S4:**
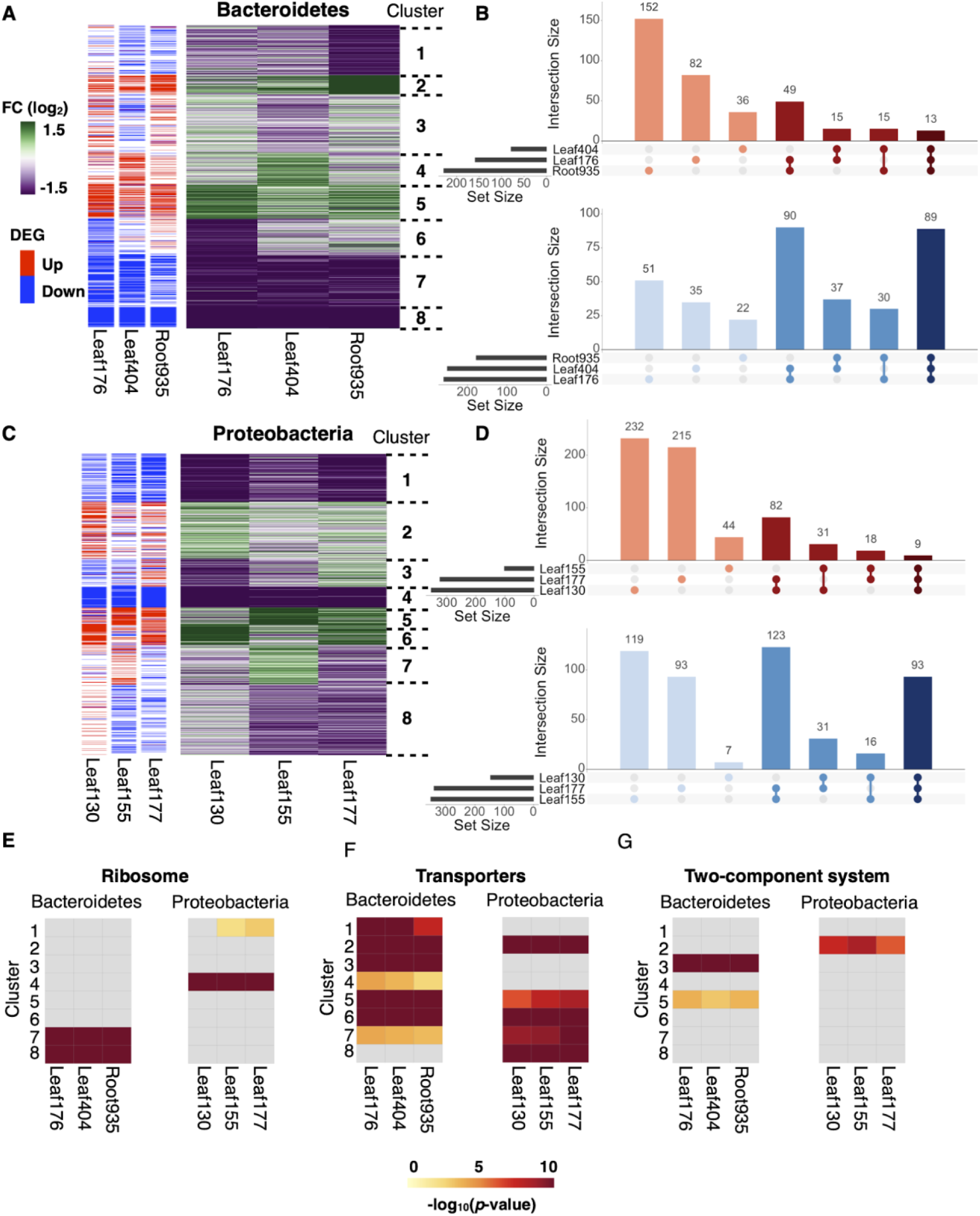
Intraphylum comparative transcriptomics of commensals. **(A and C)** Gene expression fold changes between *in planta* and *in vitro* (rich media) of **(A)** Bacteroidetes and **(C)** Proteobacteria strains. Orthologous groups shared among three strains were used for the analysis. Differentially expressed genes (DEGs) (*in planta* vs. *in vitro*; |log_2_FC| > 1; FDR < 0.01; two-tailed Student’s t test followed by Storey’s q-value) are indicated in the sidebars. Gene clusters defined by k-mean clustering are shown (k = 8). **(B and D)** UpSet intersection plots of DEGs either up- (red) or down (blue)-regulated *in planta* in the **(B)** Bacteroidetes and **(D)** Proteobacteria strains. Intersection size and set size indicate the number of shared DEGs and the number of DEGs in each strain, respectively. **(E-G)** Enrichment analysis of genes with the KEGG orthology terms **(E)** “Ribosome”, **(F)** “Transporters”, and **(G)** “Two-component system”.

**Fig. S5:**
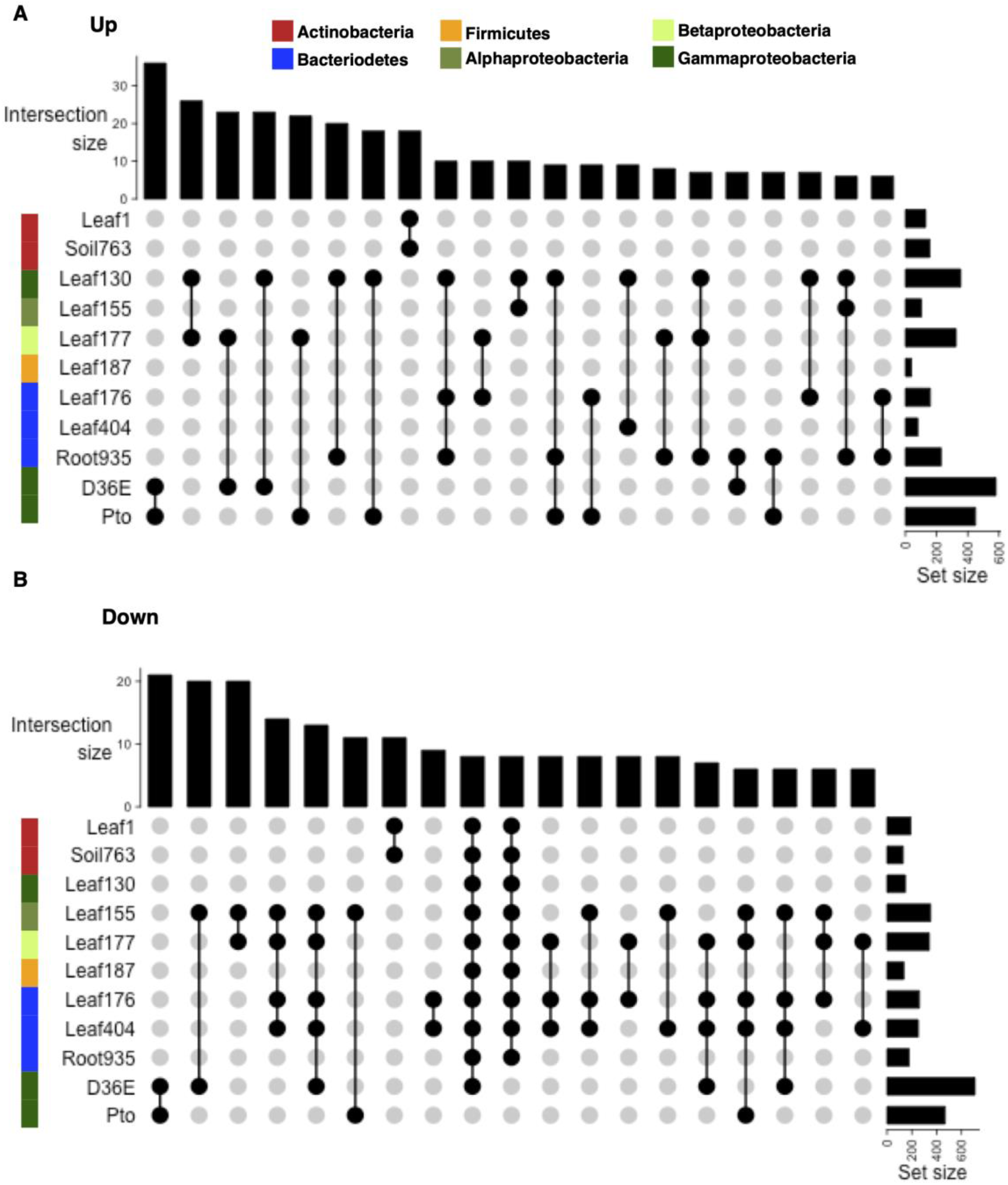
Bacterial genes differentially regulated in plants. UpSet intersection plots of differentially expressed genes (DEGs; |log_2_FC| > 1; FDR < 0.01; two-tailed Student’s t test followed by Storey’s q-value) either **(A)** up- or **(B)** downregulated *in planta*. Intersection size and set size indicate the number of shared DEGs and the number of DEGs in each strain, respectively. Combinations of more than one strain with intersection size > 5 are shown. The color sidebars indicate the taxonomic affiliation.

**Fig. S6:**
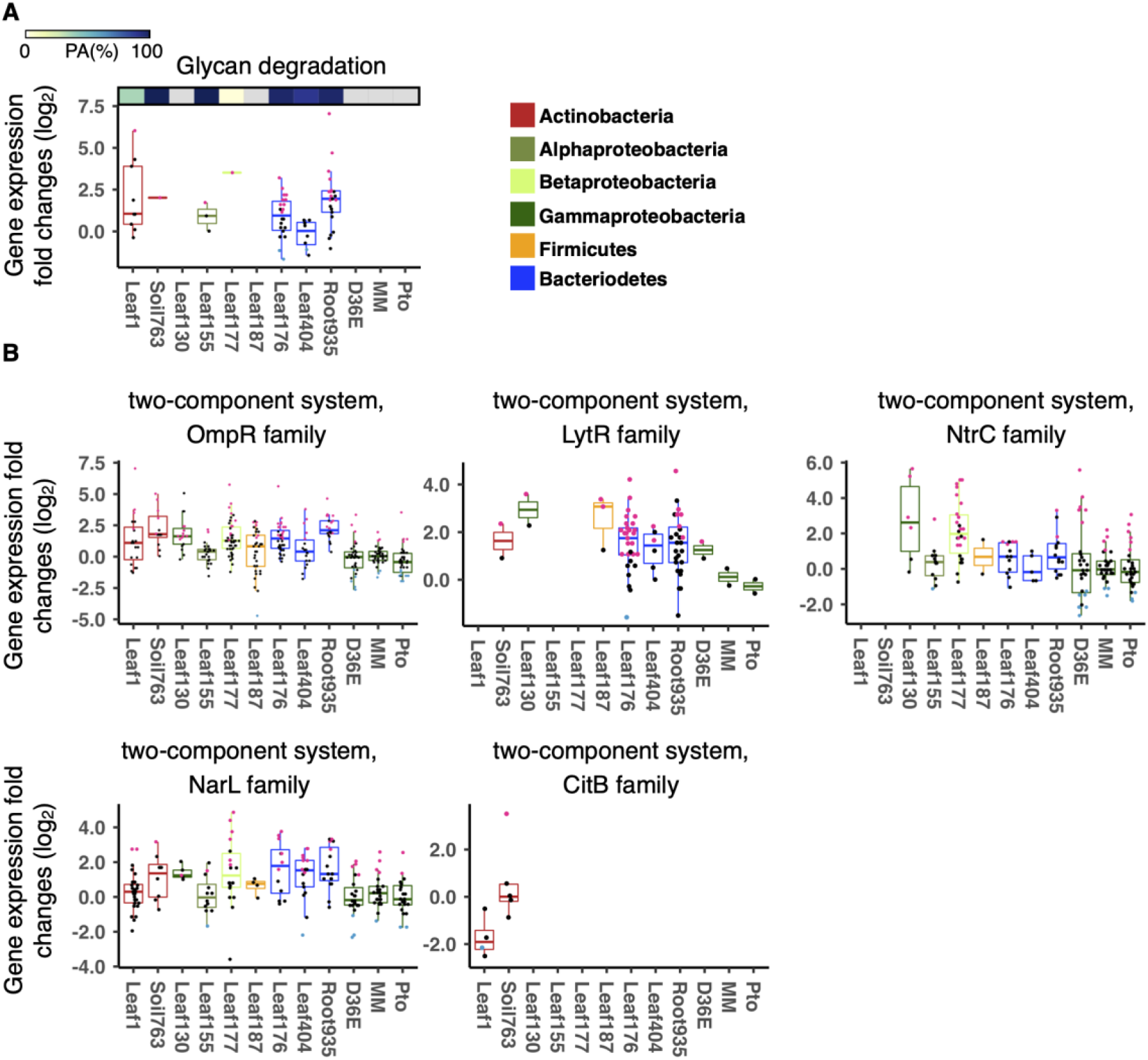
Expression of various processes of commensals *in planta*. Expression fold changes (*in planta* vs. *in vitro*) of genes related to **(A)** glycan degradation and **(B)** two-component system. Results are shown as box plots with boxes displaying the 25th–75th percentiles, the centerline indicating the median, whiskers extending to the minimum, and maximum values no further than 1.5 inter- quartile range. Box color indicates the taxonomic affiliation (phylum/class level) of each strain. All individual data points (genes) are overlaid with colors for DEGs (red: upregulated, blue: downregulated, black: non-DEG). **(A)** The top bar indicate the ratio of PA genes in each strain. All individual data points (genes) are overlaid to the box plots with colors for DEGs (red: upregulated, blue: downregulated, black: non-DEG).

**Fig. S7:**
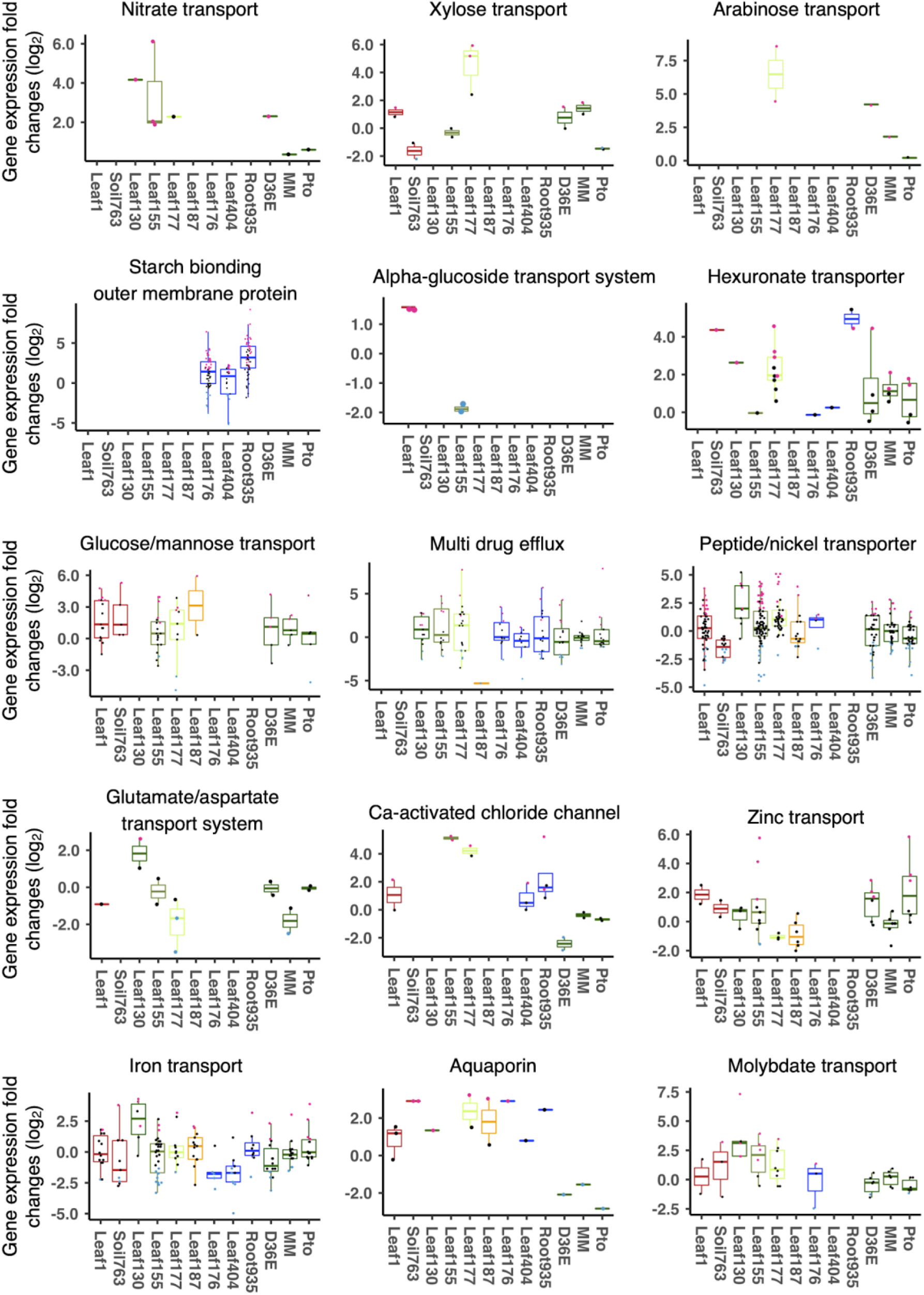
Expression of genes related to nutrient acquisition processes in commensals. Expression fold changes (*in planta* vs. *in vitro*) of genes related to nutrient transporters. Results are shown as box plots with boxes displaying the 25th–75th percentiles, the centerline indicating the median, whiskers extending to the minimum, and maximum values no further than 1.5 inter-quartile range. Box color indicates the taxonomic affiliation (phylum/class level) of each strain (see Fig. S6 for the color code). All individual data points (genes) are overlaid with colors for DEGs (red: upregulated, blue: downregulated, black: non-DEG).

**Fig. S8:**
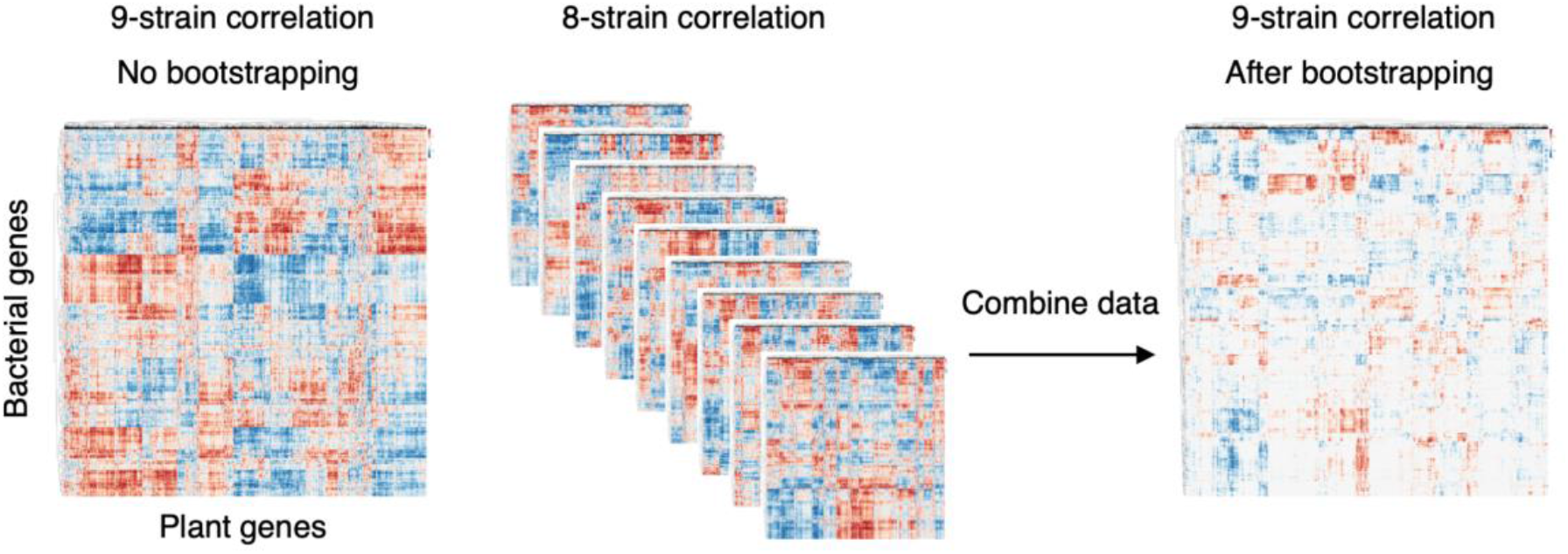
Integration of plant and bacterial transcriptomes. Schematic diagram showing a bootstrapping approach to evaluate correlations between individual plant and bacterial genes. To obtain robust correlation scores, Pearson’s correlation coefficients were calculated using all the combinations of eight strains as well as using all the nine strains. Among these 8-strain and 9-strain datasets, the weakest correlation coefficient value was used for each combination of a bacterial OG and a plant gene (“Combining data”).

**Fig. S9:**
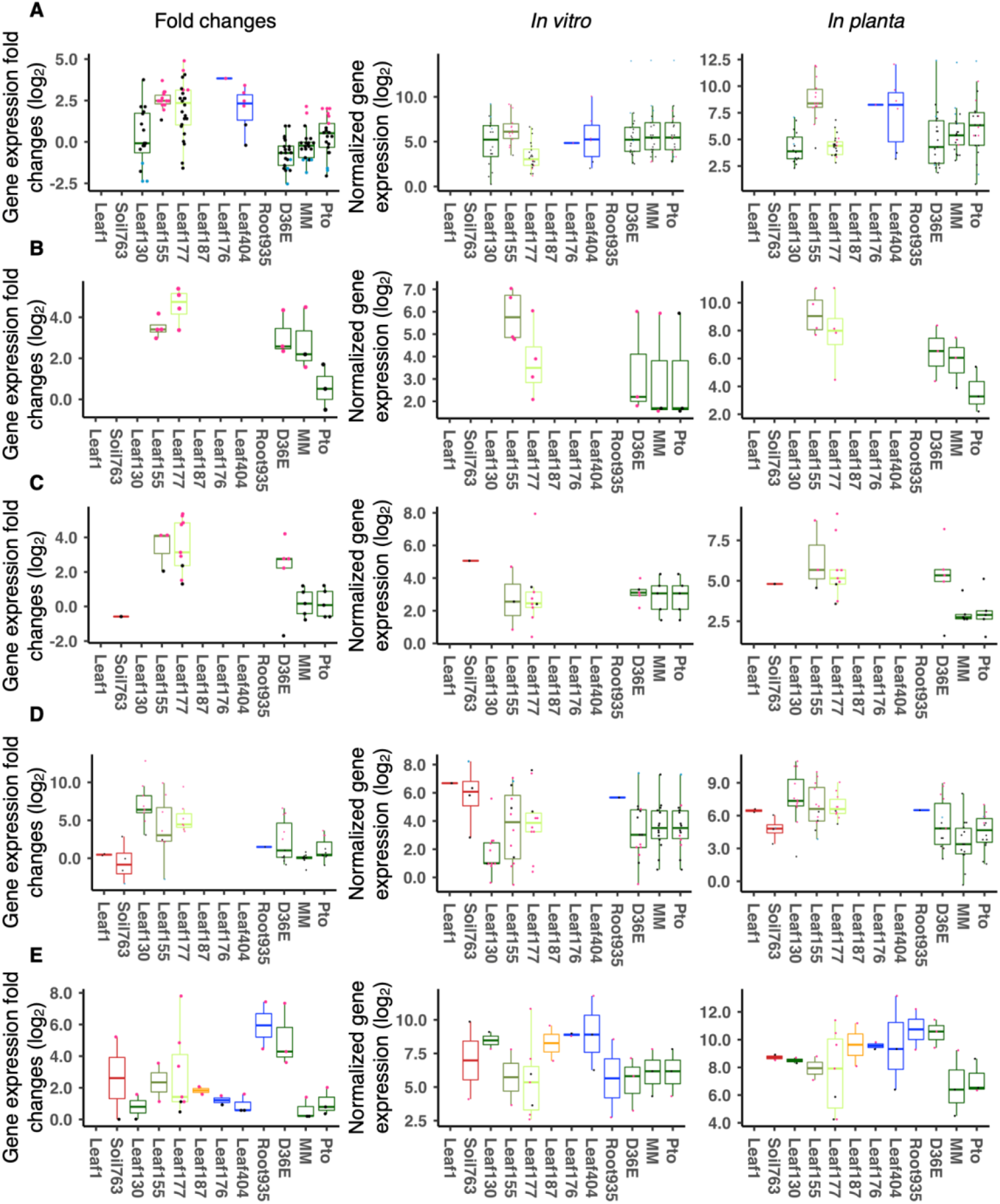
Bacterial gene expression *in vitro* and *in planta*. Expression of bacterial genes related to (A) Type 6 secretion system (B) Glycerol transport (C) Urea transport (D) Sulfur transport (E) Catalase. Results are shown as box plots with boxes displaying the 25th–75th percentiles, the centerline indicating the median, whiskers extending to the minimum, and maximum values no further than 1.5 inter- quartile range. Box color indicates the taxonomic affiliation (phylum/class level) of each strain. All individual data points (genes) are overlaid with colors for DEGs (red: upregulated, blue: downregulated, black: non-DEG).

**Fig. S10:**
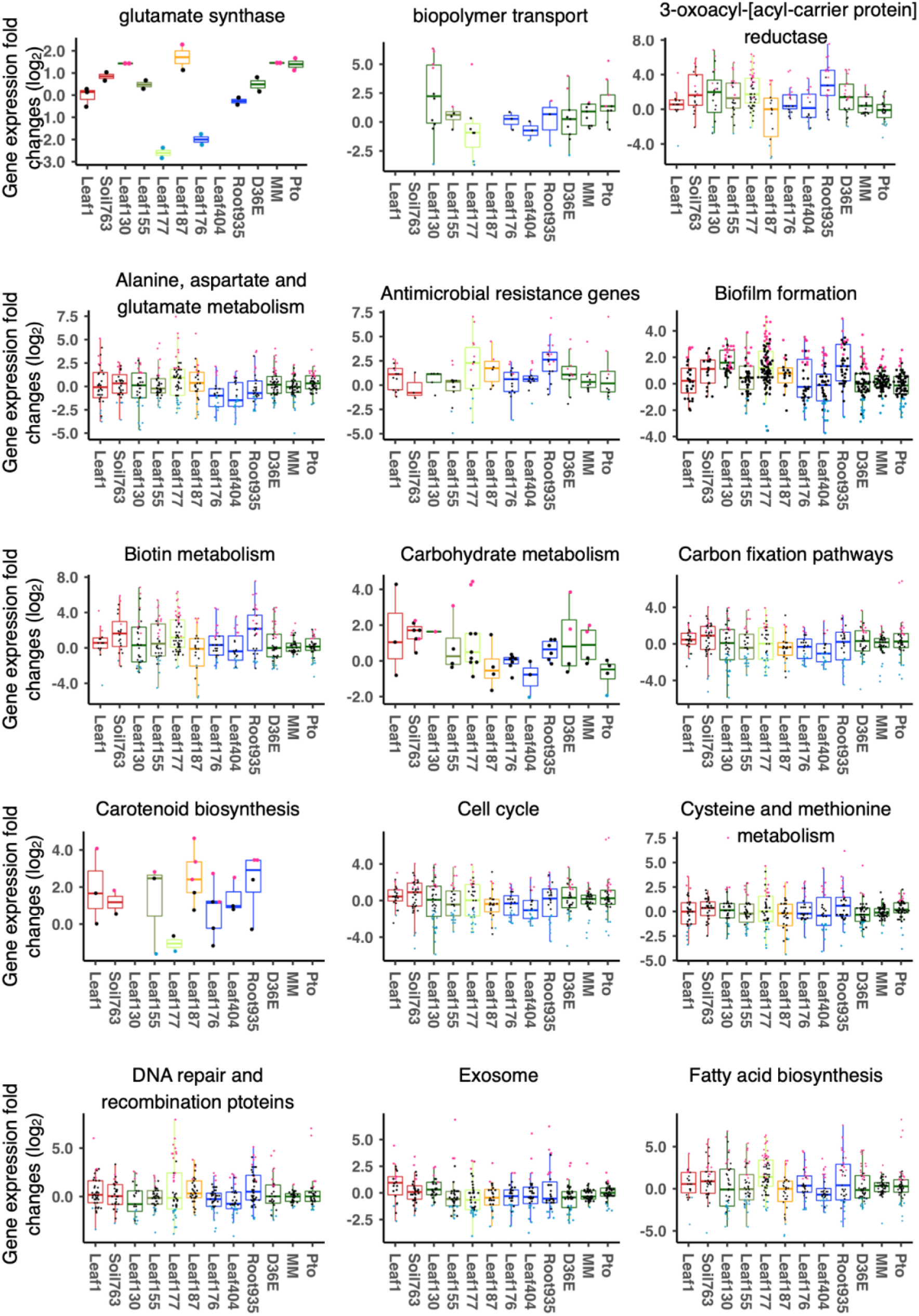
Expression of commensals genes related to various physiological processes *in planta*. Expression fold changes (*in planta* vs. *in vitro*) of genes related to various functions. Results are shown as box plots with boxes displaying the 25th–75th percentiles, the centerline indicating the median, whiskers extending to the minimum, and maximum values no further than 1.5 inter-quartile range. Box color indicates the taxonomic affiliation (phylum/class level) of each strain. All individual data points (genes) are overlaid with colors for DEGs (red: upregulated, blue: downregulated, black: non-DEG).

**Fig. S11:**
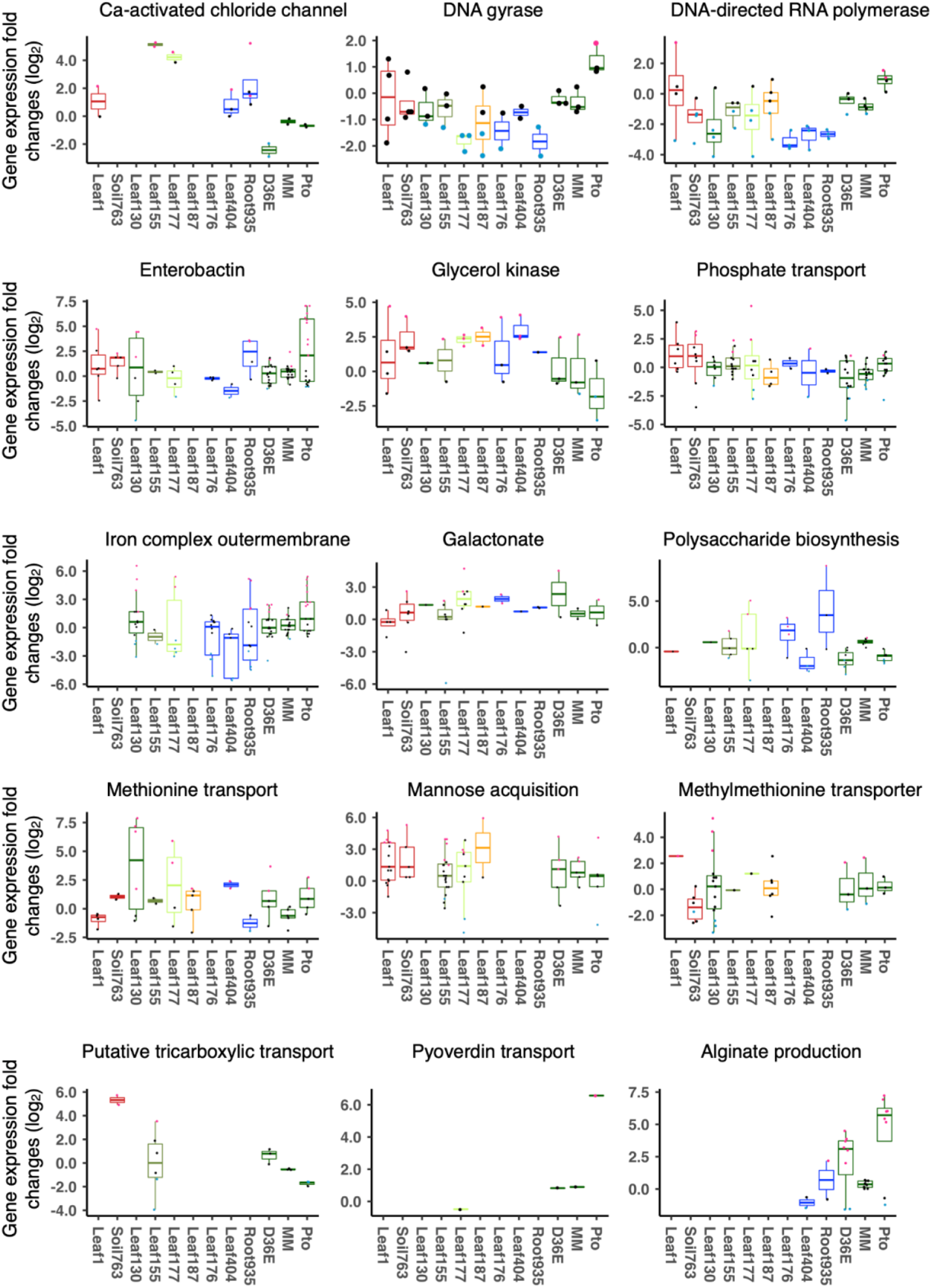
Expression of commensal genes related to various physiological processes *in planta*. Expression fold changes (*in planta* vs. *in vitro*) of genes related to various functions. Results are shown as box plots with boxes displaying the 25th–75th percentiles, the centerline indicating the median, whiskers extending to the minimum, and maximum values no further than 1.5 inter-quartile range. Box color indicates the taxonomic affiliation (phylum/class level) of each strain. All individual data points (genes) are overlaid with colors for DEGs (red: upregulated, blue: downregulated, black: non-DEG).

**Fig. S12:**
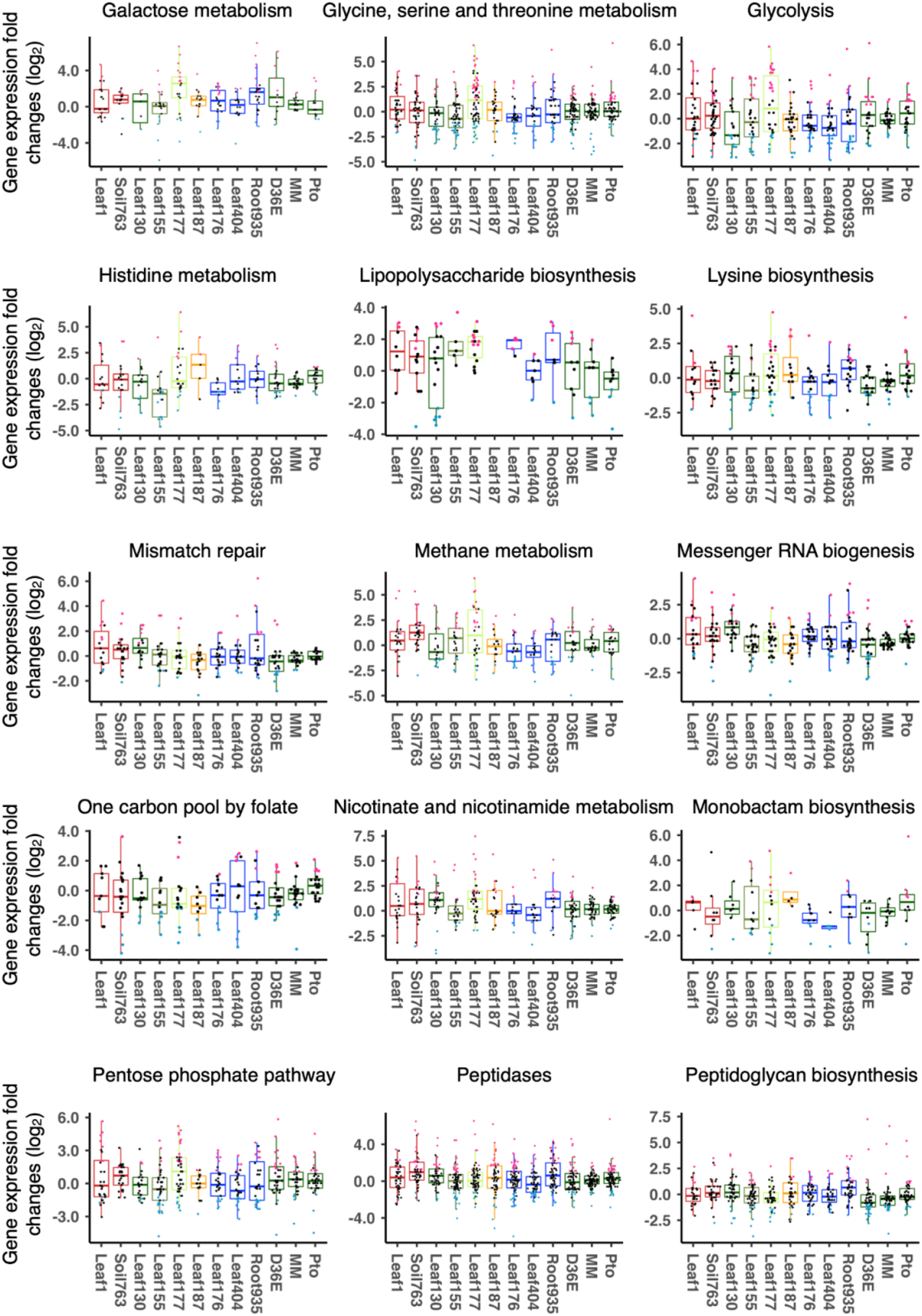
Expression of commensal genes related to various physiological processes *in planta*. Expression fold changes (*in planta* vs. *in vitro*) of genes related to various functions. Results are shown as box plots with boxes displaying the 25th–75th percentiles, the centerline indicating the median, whiskers extending to the minimum, and maximum values no further than 1.5 inter-quartile range. Box color indicates the taxonomic affiliation (phylum/class level) of each strain. All individual data points (genes) are overlaid with colors for DEGs (red: upregulated, blue: downregulated, black: non-DEG).

**Fig. S13:**
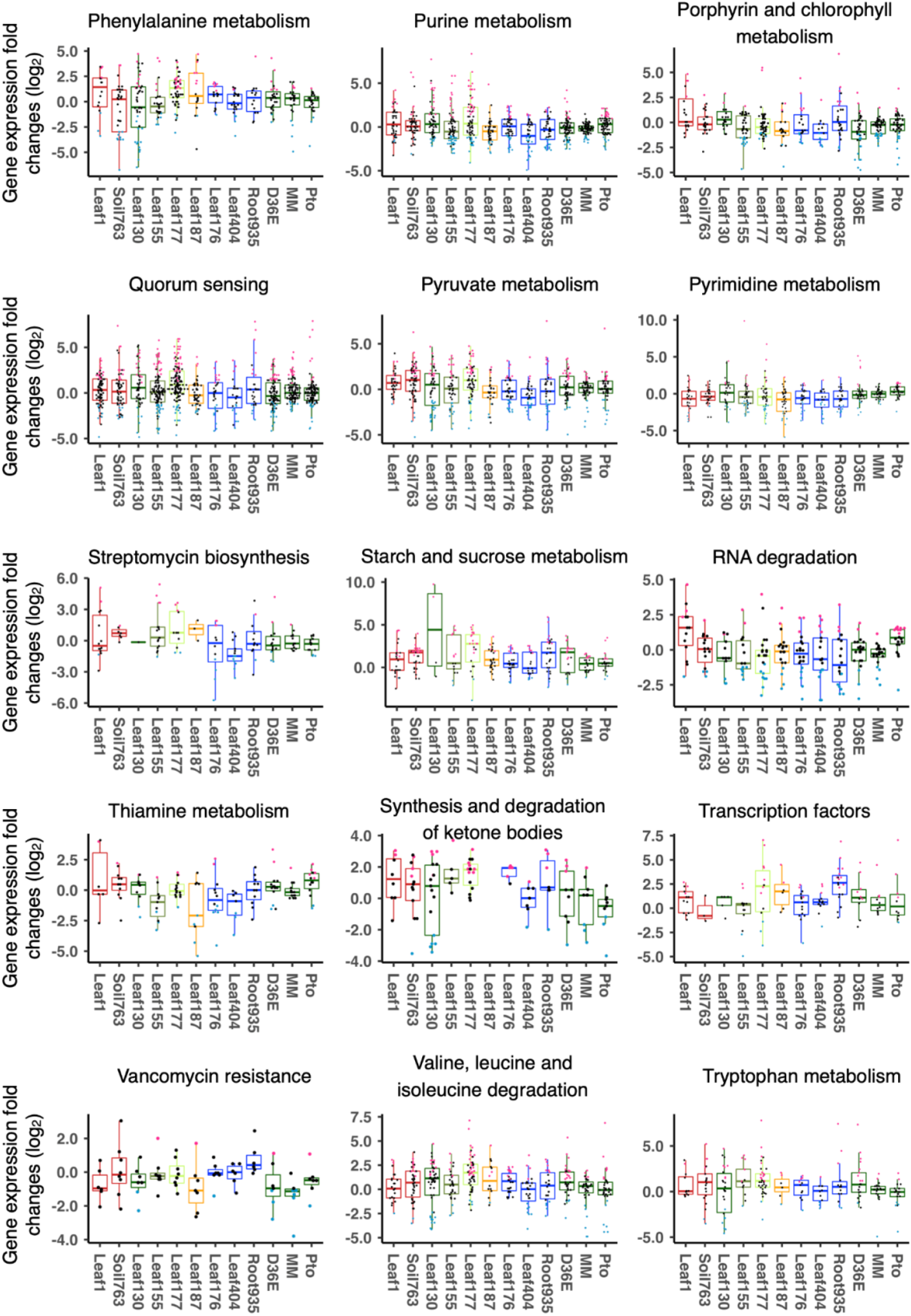
Expression of commensal genes related to various physiological processes *in planta*. Expression fold changes (*in planta* vs. *in vitro*) of genes related to various functions. Results are shown as box plots with boxes displaying the 25th–75th percentiles, the centerline indicating the median, whiskers extending to the minimum, and maximum values no further than 1.5 inter-quartile range. Box color indicates the taxonomic affiliation (phylum/class level) of each strain. All individual data points (genes) are overlaid with colors for DEGs (red: upregulated, blue: downregulated, black: non-DEG).

**Fig. S14:**
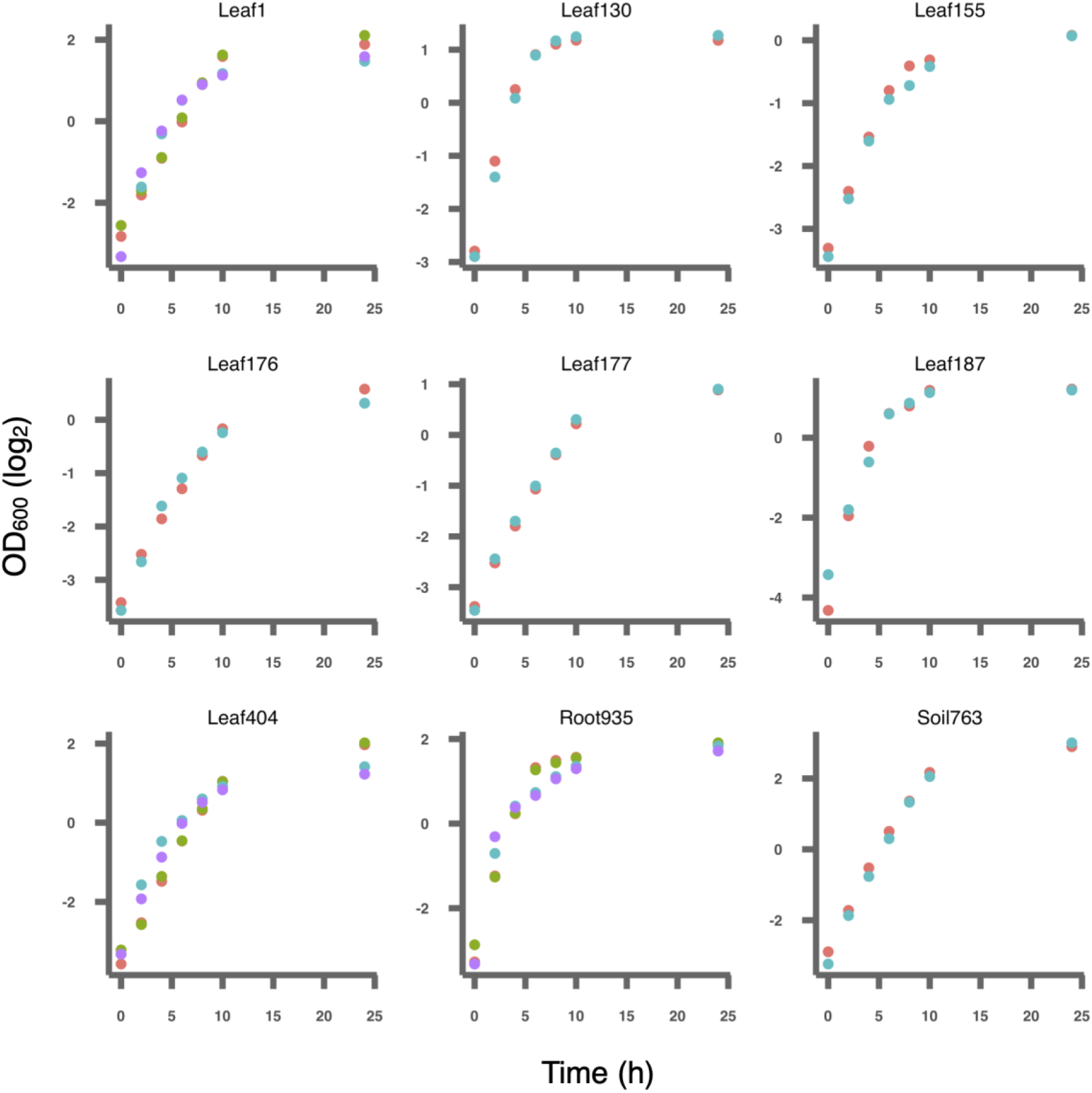
*In vitro* growth of commensals. Commensal bacteria were cultured in rich media. Replicates are shown in different colors.

